# ProtAff: Protein Binding Affinity Prediction via LoRA-Finetuned ESM-2

**DOI:** 10.64898/2026.06.13.732058

**Authors:** Lee-Shin Chu, Jeff Vogt, Michael Chungyoun, Jeffrey J. Gray

## Abstract

Predicting the binding affinity of protein–protein interactions remains a central challenge in computational biology. Structure prediction models such as AlphaFold3 (AF3) and Boltz-2 can produce high-quality docking poses, and their confidence scores indicate structure quality, but these same scores fail to rank binding affinity among confirmed binders. Here we present ProtAff, a sequence-only affinity prediction model built on ESM-2 (650M parameters) with low-rank adaptation (LoRA) fine-tuning and a cross-attention module. ProtAff is trained using a margin ranking loss on 362,567 affinity measurements spanning 20 heterogeneous data sources, and we removed all training samples whose target sequence exceeds 50% similarity to the test target EGFR. On the AdaptyvBio EGFR benchmark (*N* =55), ProtAff achieves a Spearman correlation coefficient *ρ* = 0.413, outperforming the best AF3 metric (*ρ* = 0.054), the best Boltz-2 metric (*ρ* = −0.046), and ML-based predictors MINT (*ρ* = 0.242) and CrossAffinity (*ρ* = 0.216). Applied to the AdaptyvBio Nipah virus binder design competition, a pipeline incorporating ProtAff for affinity ranking produced a design with *K*_D_ = 0.132 nM (2 of 5 designs confirmed binding), a 2.8-fold improvement over the competition winner. On a cross-target discrimination benchmark of 91 VHH–antigen crystal structures, ProtAff underperforms structural methods for distinguishing cognate from non-cognate pairings, indicating that sequence-based affinity models are effective for within-target ranking but not for cross-target specificity.

## 1 Introduction

Quantifying the strength of protein–protein binding, typically expressed as the dissociation constant *K_D_*, is essential both for understanding biological mechanisms and for engineering therapeutic proteins such as antibodies. Although deep learning has dramatically improved protein complex structure prediction, accurately predicting binding affinity from sequence or structure remains an open problem. Models such as AlphaFold3 (AF3) [1], Boltz-2 [2], and Chai-1 [3] predict protein complex structures with high accuracy, and their confidence scores (ipTM, ipSAE [4], pLDDT) have been widely adopted as computational filters in protein design pipelines. These scores perform well for assessing structural quality: Smorodina et al. [5] benchmarked AF3, Boltz-2, and Chai-1 on 106 experimentally determined nanobody–antigen complexes and found that AF3’s ipTM correlates strongly with docking quality (DockQ), achieving Pearson correlation coefficient *r* = 0.938. They are also effective for binary binder classification: Overath et al. [6] meta-analyzed 3,766 experimentally tested de novo designs across 15 targets and confirmed that AF3 ipSAE is the most predictive structural metric for distinguishing binders from non-binders, with a 1.4-fold improvement in average precision over ipAE.

However, classifying binders from non-binders or ranking docking pose quality does not translate into predicting binding affinity. Binder selection is a binary classification task (distinguishing protein partners that bind from those that do not), whereas affinity prediction requires ranking the binding strength among confirmed binders. A structural metric can be effective at the former by rejecting misfolded or non-interacting designs, yet still fail at the latter, because confidence scores capture interface plausibility, not binding energetics. On the AdaptyvBio EGFR benchmark of 55 single-chain protein binders with experimentally measured −log *K_D_* values [7], all structural metrics from AF3 and Boltz-2 showed negative or near-zero Spearman correlations with affinity (Section 4.1).

The gap between structural pose quality and binding affinity prediction motivates the development of dedicated affinity models. Traditionally, affinity prediction relied on physics-based energy functions such as Rosetta flex ddG [8] and FoldX [9], or on contact-based statistical models such as PRODIGY [10]. These methods require experimentally determined or accurately predicted complex structures as input, limiting their applicability to novel sequences without experimental structures. A growing body of machine learning work has since targeted protein–protein affinity prediction using sequence-based, structure-based, and multimodal approaches [11, 12]. However, existing methods face several limitations. Hummer et al. [13] found that an equivariant graph neural network for antibody-antigen ΔΔ*G* prediction achieves high accuracy under random splits but degrades under complementarity-determining region (CDR)-sequence-identity cutoffs. The AbRank benchmark [14] shows high sequence similarity between training and test splits, which is relevant when interpreting reported performance (Figure S1). More broadly, most methods are evaluated on within-distribution splits [15] or focus on ΔΔ*G* prediction for point mutations, leaving generalization across novel targets and heterogeneous assay types unclear.

Among recent PLM-based approaches, PLM-interact [16], AntiBinder [17], and PPLM [18] jointly encode protein pairs for binary interaction prediction but not for continuous affinity ranking. AlphaBind [19] pre-trains on millions of binding measurements for guided affinity optimization, and several groups have explored parameter-efficient fine-tuning of PLMs for antibody fitness tasks [20, 21]. Most directly comparable to our work, MINT [22] is pre-trained on unlabeled paired protein sequences from the STRING database and fine-tuned for binding affinity prediction. CrossAffinity [23] employs cross-attention between binder and target representations, sharing architectural similarities with our approach but without fine-tuning the PLMs. At the same time, protein language models have shown the capacity to capture biophysical information from sequence alone, motivating a dedicated sequence-based framework trained on large-scale, heterogeneous binding data with rigorous train-test separation.

In this work, we present ProtAff, a sequence-only binding affinity prediction model. Our contribution lies in the novel integration of LoRA fine-tuning, cross-attention, and margin ranking loss into a unified framework trained on a large-scale, curated dataset. Specifically: (1) we construct a training set of 362,567 affinity measurements spanning 20 heterogeneous data sources, filtered to ensure no training target exceeds 50% sequence similarity to the test target; (2) we demonstrate that ProtAff outperforms both structural confidence scores and existing PLM-based affinity predictors on the AdaptyvBio EGFR benchmark; (3) we validate practical utility in a binder design pipeline that produced a 2.8-fold stronger binder than the winner of the AdaptyvBio Nipah virus glycoprotein G design competition; and (4) we characterize where sequence-only affinity models succeed and fail through cross-target discrimination, target ablation, and biophysical property analyses, showing that structural methods retain advantages for cross-target specificity and that ProtAff’s within-target ranking correlates with binder-level compositional features rather than target-conditioned interaction modeling.

## 2 Methods

### 2.1 Data Curation

Each affinity measurement in our dataset describes a pair of interacting proteins: a *binder* (the protein whose binding strength is being assessed, typically an antibody or a computationally designed protein) and a *target* (the protein it binds to, such as a viral surface protein or a cell-surface receptor). We assembled a training set of 362,567 affinity measurements by integrating data from three aggregated databases: AbRank [14], ASD [24], and PPB-Affinity [25]. Together, these sources compile affinity data from 20 original repositories [26–43] (Figure 1A; Table S1) and include 308,854 unique binders to 10,949 unique targets. The raw data span a wide range of assay types (*K_D_*, IC_50_, and escape ratios from deep mutational scanning), covering more than 15 orders of magnitude in reported values. Following AbRank [14], all measurements are approximately converted to a common scale of log_10_(*K_D_* [nM]): *K_D_* values are converted directly, IC_50_ values are treated as approximate *K_D_* estimates, and escape ratios are log-transformed and linearly rescaled to the log_10_(*K_D_*) range. These conversions are approximate, and residual scale differences between assay types remain; the margin ranking loss (Section 2.3) mitigates this by training on ordinal relationships rather than absolute values.

**Figure 1:**
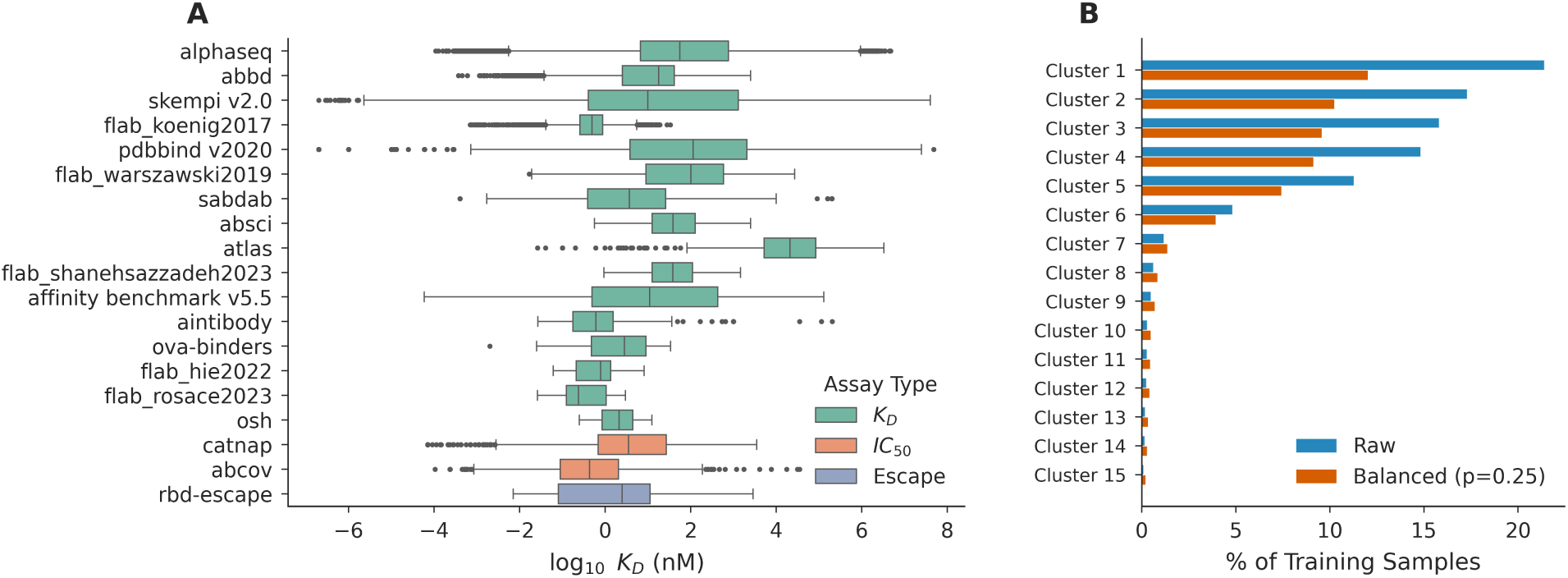
Overview of the training dataset. **(A)** Distribution of log_10_ *K_D_*(nM) across 20 data sources (362,567 samples total), colored by assay type (*K_D_*, *IC*_50_, escape) and sorted by sample count within each assay group. **(B)** Raw representation (blue) and effective sampling weight after power-law balancing (*p* = 0.25, orange) for the top 15 clusters by size.

The training data exhibit severe target imbalance: five targets account for more than 280,000 of the 362,567 total samples, while most of the 10,949 unique targets have fewer than 100 associated measurements (Figure 1B). Clustering all target sequences at 90% sequence identity using MMseqs2 [44] yields 3,158 clusters with a highly skewed distribution; a single cluster contains approximately 77,000 samples (21% of the dataset). The five largest clusters are each dominated by a single target: SARS-CoV-2 Spike ectodomain (77,579 samples, 1273 amino acids (aa)), human PD-1 (62,720 samples, 123 aa), human TIGIT (57,304 samples, 116 aa), SARS-CoV-2 RBD (a subdomain of Spike, 53,735 samples, 209 aa), and influenza hemagglutinin H1 (40,878 samples, 519 aa).

### 2.2 Model Architecture

ProtAff builds on ESM-2 (650M parameters) [45], a protein language model pretrained on large-scale protein sequence databases using masked language modeling. A shared ESM-2 backbone encodes both the binder and target sequences independently (Figure 2). We apply low-rank adaptation (LoRA) [46] to the query, key, and value projections of the last four transformer layers. For each adapted weight matrix *W*_0_ ∈ R*^d^*^×*d*^, LoRA introduces two low-rank matrices such that the modified forward pass computes *W*_0_*x* + *BAx*, where *A* ∈ R*^r^*^×*d*^ and *B* ∈ R*^d^*^×*r*^ with rank *r* = 16 and scaling factor *α* = 32 (which controls the magnitude of the adapter update relative to the pretrained weights). The matrix *A* is initialized from *N* (0, σ ^2^) and *B* is initialized to zero, so the adapter output is zero at the start of training and the pretrained weights are unperturbed. The original weights *W*_0_ remain frozen throughout training, and only the LoRA parameters are updated. This enables efficient task-specific adaptation while preserving the pretrained representations, an approach shown to achieve competitive performance with full fine-tuning for protein interaction tasks [47].

**Figure 2:**
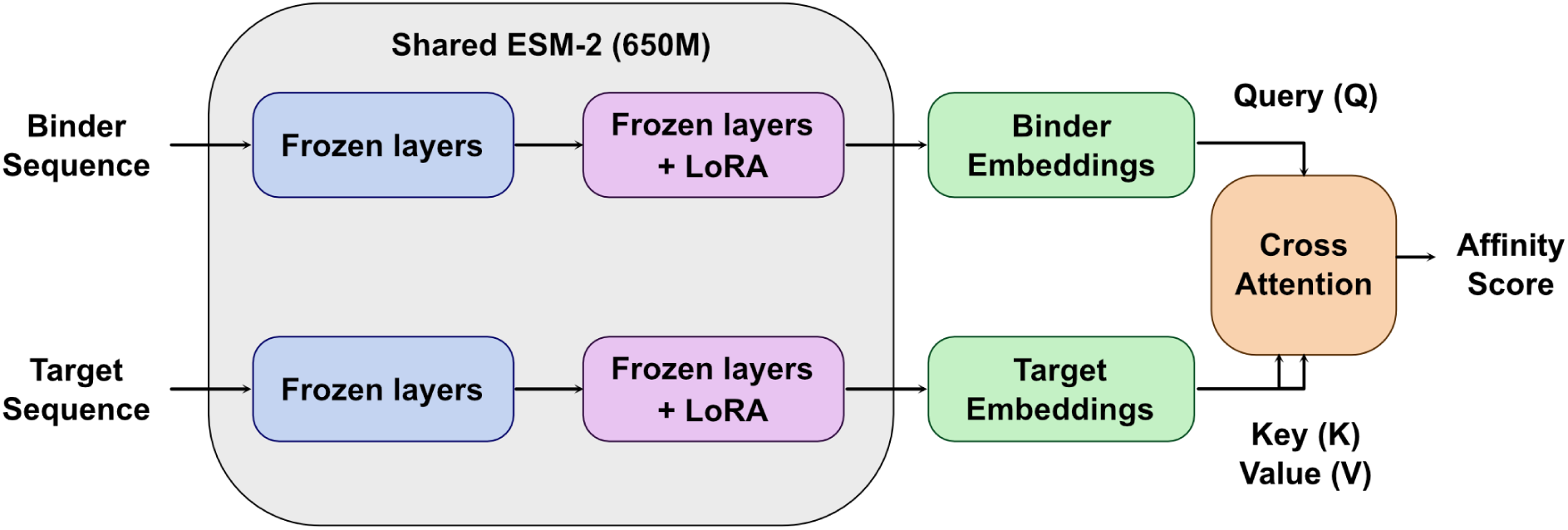
Overview of the ProtAff architecture. Binder and target sequences are independently encoded by a shared ESM-2 (650M) backbone, where early layers are frozen and later layers are frozen with LoRA adapters. The resulting binder embeddings serve as queries (Q) and target embeddings as keys (K) and values (V) in a cross-attention module, which produces a scalar affinity score.

After the finetuned ESM-2, the 1280-dimensional per-residue embeddings are projected to *d*_model_ = 256 dimensions via a linear layer with layer normalization. The binder embeddings then serve as queries and the target embeddings as keys and values in a two-layer cross-attention module [48] with 8 heads per layer and scaled dot-product attention, consistent with recent findings that cross-attention outperforms naive concatenation of protein representations for protein-protein interaction (PPI) tasks [49]. The unidirectional formulation reflects the asymmetry of the design task: for a fixed target, the goal is to rank different binders by their binding strength, so the binder representation queries the target rather than the reverse. The cross-attention output is passed through a learned attention pooling layer (a single-head attention mechanism that computes per-residue importance weights and produces a fixed-length representation), followed by a two-layer multi-layer perceptron (MLP) head that outputs a single scalar affinity prediction.

### 2.3 Training Objective

Because different assay types report values on incomparable scales even after the approximate conversion described in Section 2.1, we adopt a margin ranking loss that requires only ordinal correctness:

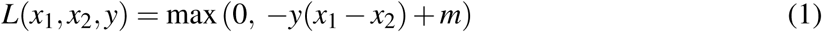

where *x*_1_ and *x*_2_ are the predicted affinity scores for two binder–target pairs, *y* ∈ {−1, +1} indicates which pair has stronger binding, and *m* = 0.1 is the margin hyperparameter. AbRank [14] previously applied margin ranking loss to antibody affinity prediction across heterogeneous assay types, demonstrating its suitability for this setting. We also evaluate a weighted variant of the margin ranking loss that scales each pair’s contribution by the magnitude of the affinity difference between the two binders:

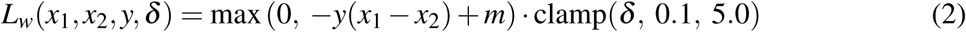

where *δ* is the absolute difference in converted log_10_(*K_D_* [nM]) between the two binders, clamped to the range [0.1, 5.0]. Pairs separated by a large affinity gap receive proportionally higher weight, encouraging the model to prioritize correct ordering of binders with meaningfully different binding strengths over fine-grained discrimination among binders with similar affinities.

### 2.4 Pair Construction and Sampling

To encourage both within-target discrimination and a globally coherent affinity scale, training pairs are constructed in two complementary regimes. Intra-target pairs compare two binders measured against the same target. For each target, binders are sorted by log_10_(*K_D_*[nM]), and pairs are formed between binders whose affinity difference exceeds a minimum gap *δ*_min_ = 1.0 (corresponding to a 10-fold difference in *K_D_*), ensuring the model learns from meaningfully distinct binding differences rather than noisy near-ties. For each anchor binder, up to two partners are sampled from the valid margin window. The number of anchor binders per target is capped at 2,000, selected by uniform random sampling when a target has more associated binders.

Inter-target pairs compare binders measured against different targets. The primary motivation is coverage: many targets in the training set have very few associated binders, and singleton targets (those with only one measured binder) cannot form any intra-target pairs. Without inter-target pairing, these samples would be excluded from training despite containing valid affinity signal. Inter-target pairing also exposes the cross-attention module to a wider variety of target sequences during training, which may help it learn more general target representations rather than overfitting to the well-represented targets that dominate intra-target pairing.

To construct inter-target pairs, two binders are sampled from each target, producing a pool of records spanning all targets. These records are sorted globally by log_10_(*K_D_* [nM]), and pairs are formed between binders bound to different targets whose affinity difference exceeds the minimum gap *δ*_min_, with up to 10 partners per anchor. Some inter-target ordinal labels are inevitably noisy because they may span different assay types, but these pairs constitute a minority of training data and serve primarily to ensure broad target coverage.

To address the severe target imbalance described in Figure 1, we employ cluster-balanced pair sampling. All target sequences are clustered at 90% sequence identity using MMseqs2. Because each training pair involves two targets, the pair-level sampling weight is derived from the geometric mean of the two cluster sizes:

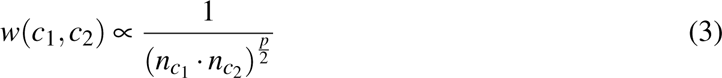

where *n_c_* is the number of samples in cluster *c* and *p* = 0.25 is the cluster balance power (Supplementary Table S2). This scaling reduces the largest cluster’s effective share from 21.4% to 12.0%.

Validation uses a group-based split in which all validation targets are entirely unseen clusters during training, providing a realistic estimate of generalization.

### 2.5 Hyperparameters

All sequences are truncated to a maximum of 1,024 residues. Training uses the AdamW optimizer with linear warmup over the first 10% of steps, gradient clipping at norm 1.0, and bf16 mixed precision. Full hyperparameters are listed in Supplementary Table S2.

## 3 Experiments

### 3.1 Affinity Ranking on AdaptyvBio EGFR Benchmark

We evaluated ProtAff on the AdaptyvBio EGFR benchmark [7] (*N* =55), consisting of single-chain protein binders under 250 residues with experimentally measured dissociation constants against EGFR. The test target (EGFR) shares *<*50% sequence similarity with any target in the training data, making it a stringent test of generalization (Figure 3).

**Figure 3:**
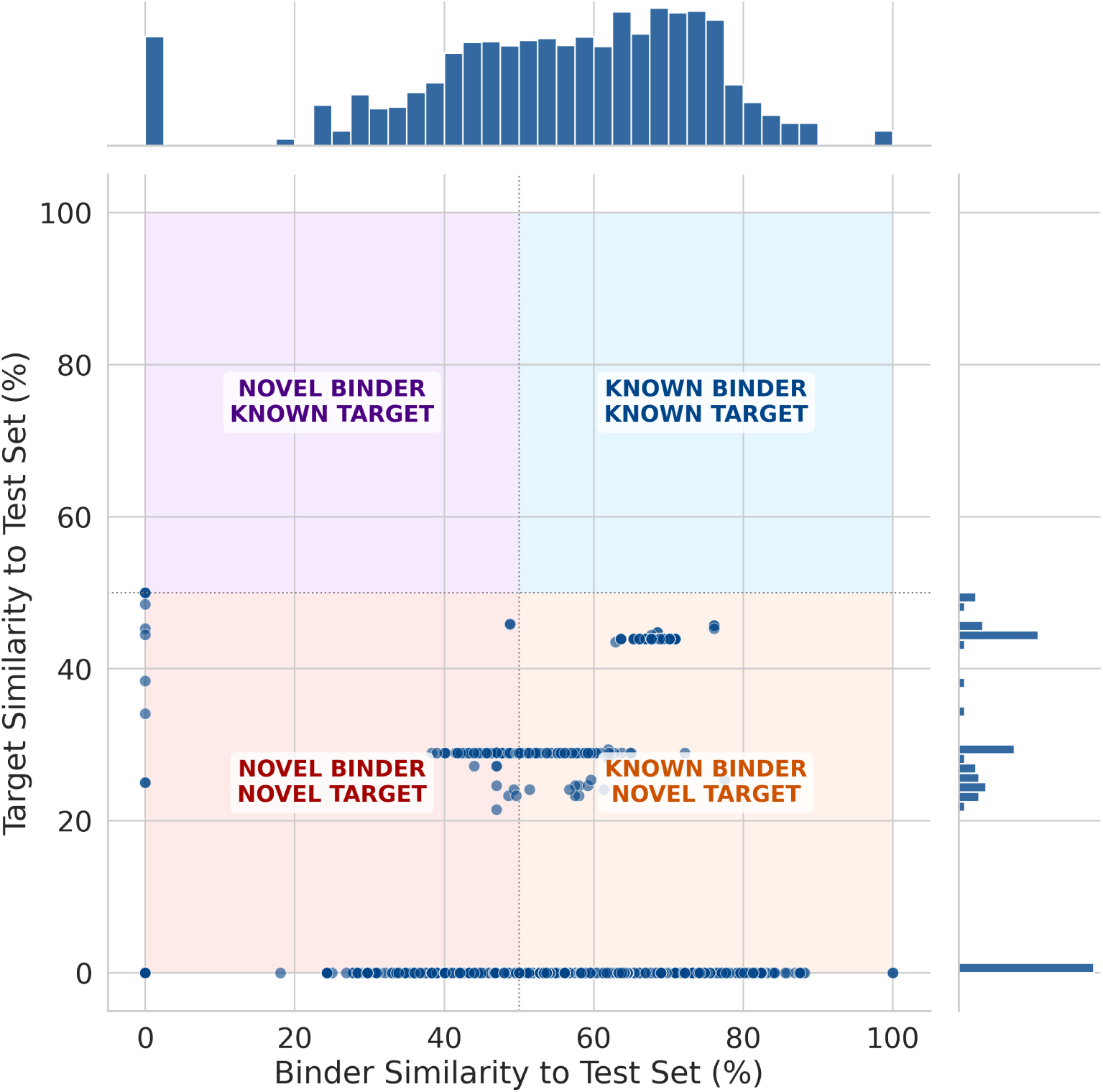
Training set sequence similarity to the AdaptyvBio EGFR benchmark (*N* = 55). Each point represents a training pair, plotted by maximum binder sequence similarity (*x*-axis) and maximum target sequence similarity (*y*-axis) to the test set.

We compared ProtAff against structural confidence metrics from both Boltz-2 [2] and AF3 [1] (ipSAE [4], ipTM, pDockQ [50], pDockQ2 [51], and LIS [52]), each aggregated across multiple seeds using min, max, mean, and median (Supplementary Table S3), as well as two ML-based affinity predictors: MINT [22] and CrossAffinity [23]. We report Spearman rank correlation (*ρ*) and Pearson correlation (*R*) between predicted scores and experimental −log *K_D_* for overall ranking quality, and normalized discounted cumulative gain at *K* (NDCG@*K*) [53] for top-of-list ranking accuracy. NDCG@*K* uses continuous −log *K_D_* values as graded relevance scores and applies logarithmic discounting to penalize misordering at higher ranks more heavily. Specifically, NDCG@*K* is defined as

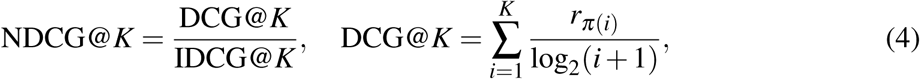

where *π*(*i*) is the index of the item placed at rank *i* by the predicted scores, *r_π_*_(*i*)_ is the graded relevance (here, −log *K_D_*) of the item ranked at position *i*, and IDCG@*K* is the DCG@*K* obtained under the ideal (ground-truth) ranking. This metric better reflects practical screening scenarios where only a small number of candidates are selected for experimental validation.

### 3.2 Application: AdaptyvBio Nipah Virus Binder Design Competition

To assess ProtAff in a prospective design setting, we designed binders for Nipah virus glycoprotein G (PDB: 2VSM) as part of the AdaptyvBio binder design competition held in Nov. 2025 [54]. Our designs were not selected for the official competition round but were experimentally tested at the same time by the competition organizers on the same platform using assay credits, ensuring comparable experimental conditions.

We employed a three-stage pipeline starting from the heavy and light chains of the 1E5 Fab antibody [55] (PDB: 8K0C) (Supplementary Figure S2). In Stage 1 (sampling), we used AntiB-ERTy [56] as a masked language model to identify mutable positions (those where the wild-type residue has low pseudo-log-likelihood) and stochastically sampled mutation combinations to generate 10,000 candidate variants, each with 10 mutations away from the 1E5 Fab. In Stage 2 (filtering), we retained the top 100 candidates by combined AntiBERTy pseudo-log-likelihood and IgLM [57] perplexity scores, averaged over heavy and light chains. In Stage 3 (affinity ranking), we scored the remaining candidates against Nipah glycoprotein G using ProtAff and selected the five designs with the highest predicted affinity. AdaptyvBio used biolayer interferometry (BLI) to measure *K_D_* experimentally.

## 4 Results

### 4.1 ProtAff Outperforms Structural and ML-Based Baselines for Within-Target Affinity Ranking

On the AdaptyvBio EGFR benchmark, ProtAff achieved a Spearman *ρ* = 0.413 and the margin-weighted variant achieved *ρ* = 0.381 between predicted affinity and experimental −log *K_D_* (Figure 4). We report AF3 structural metrics using min aggregation across seeds, which selects the most conservative confidence estimate per design and yielded the highest Spearman *ρ* among the four aggregation strategies tested (Supplementary Table S3). All AF3 structural metrics showed negative or near-zero correlations with affinity, with the best (ipSAE, min aggregation) marginally positive but statistically indistinguishable from zero. The remaining metrics (ipTM, pDockQ, pDockQ2, LIS) were all negative (Table 1). Boltz-2 structural metrics performed even worse, with uniformly negative correlations (Supplementary Table S3). Two ML-based affinity predictors, MINT [22] (*ρ* = 0.242) and CrossAffinity [23] (*ρ* = 0.216), both outperform structural metrics but fall short of ProtAff across all evaluation metrics (Table 1).

**Figure 4:**
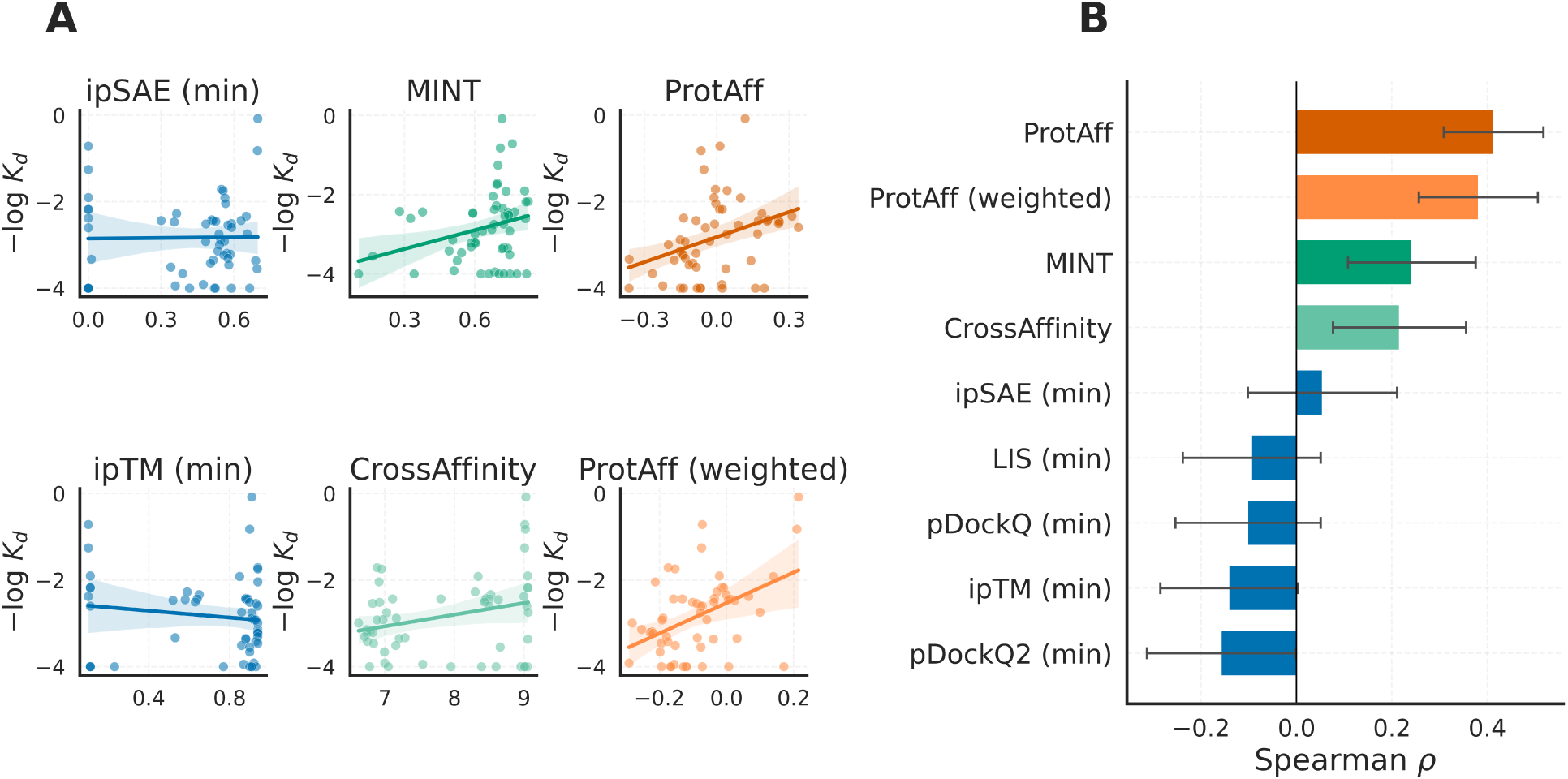
Comparison of binding affinity prediction methods on the AdaptyvBio EGFR benchmark (*N* = 55). **(A)** Scatter plots of predicted scores versus experimental −log *K_D_* for AF3 structural metrics (min aggregation), ML-based predictors (MINT, CrossAffinity), and both ProtAff configurations. Regression lines with 95% confidence intervals are shown. **(B)** Spearman rank correlation (*ρ*) for each method. Error bars indicate ± standard deviation over 1,000 bootstrap resamples.

**Table 1:** Quantitative comparison of binding affinity prediction methods on the AdaptyvBio EGFR benchmark (*N* = 55). Structural metrics are computed from AF3 docking models using min aggregation. ProtAff ablation variants (No LoRA, MSE loss) are included below the main configurations. Values are reported as mean ± std over 1,000 bootstrap resamples. Best values are in **bold**.

| Method | Spearman $\rho$ | Pearson $R$ | NDCG@5 |
| --- | --- | --- | --- |
| ipSAE (min) | 0.054±0.157 | 0.013±0.168 | 0.666±0.208 |
| ipTM (min) | -0.141±0.149 | -0.137±0.162 | 0.588±0.110 |
| pDockQ (min) | -0.102±0.156 | -0.083±0.169 | 0.274±0.088 |
| pDockQ2 (min) | -0.158±0.159 | -0.098±0.160 | 0.694±0.184 |
| LIS (min) | -0.094±0.149 | -0.076±0.158 | 0.265±0.061 |
| MINT | 0.242±0.134 | 0.275±0.103 | 0.430±0.135 |
| CrossAffinity | 0.216±0.139 | 0.277±0.134 | 0.432±0.106 |
| **ProtAff** | **0.413±0.105** | 0.343±0.087 | 0.463±0.072 |
| **ProtAff (weighted)** | 0.381±0.125 | **0.451±0.145** | **0.758±0.185** |
| No LoRA | 0.257±0.132 | 0.260±0.099 | 0.482±0.099 |
| MSE loss | 0.163±0.147 | 0.081±0.135 | 0.218±0.132 |

To isolate the contributions of individual design choices, we evaluated two ablation variants (Table 1). Replacing the margin ranking loss with mean squared error (MSE) regression on log-transformed affinity values reduces Spearman *ρ* from 0.413 to 0.163. Freezing the ESM-2 backbone entirely, removing LoRA adapters while retaining the cross-attention module and margin ranking loss, reduces *ρ* to 0.257. Both components contribute to ProtAff’s performance, with the ranking loss providing the larger individual improvement.

We also evaluated top-of-list ranking quality using NDCG@*K* across nine values of *K* between 1 and 50, reflecting practical screening settings where only a small number of candidates are selected for experimental validation. ProtAff (weighted) achieves the highest NDCG@*K* across nearly all values of *K*, outperforming all structural metrics and ML-based baselines (Figure 5). At *K* = 1, both ProtAff (weighted) and ipSAE (min) achieve perfect ranking. As *K* increases, structural metrics decline sharply while ProtAff (weighted) maintains NDCG above 0.7 through *K* = 50. The default ProtAff (margin ranking loss) starts lower at small *K* but converges with the ML baselines by *K* = 15. The two ProtAff configurations diverge most at small *K*, with the weighted variant achieving higher NDCG@*K* for *K* ≤ 10 but lower Spearman *ρ* over the full ranking (Table 1). The nature of this tradeoff is examined in Section 5.2.

**Figure 5:**
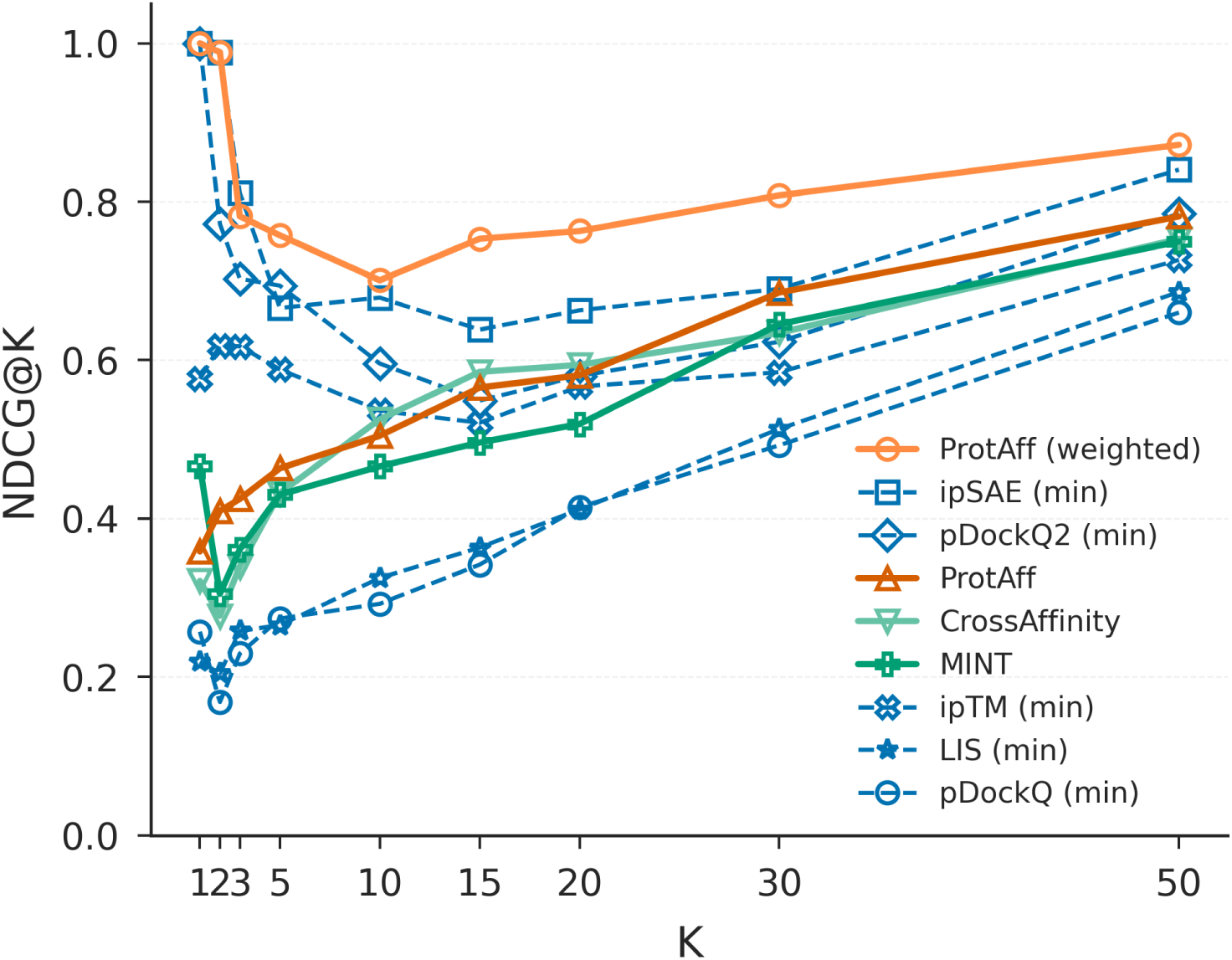
NDCG@*K* on the AdaptyvBio EGFR benchmark (*N* = 55). Solid orange lines: ProtAff variants; solid green lines: ML baselines (MINT, CrossAffinity); dashed blue lines: AF3 structural metrics (min aggregation).

### 4.2 ProtAff-Guided Design Yields a Sub-Nanomolar Nipah Virus Binder

In the AdaptyvBio Nipah virus binder design competition, our top-ranked design (Rank 1) was experimentally confirmed as a strong binder with *K_D_* = 1.32 × 10^−10^ M (131.6 pM), high expression, and three BLI replicates with a standard deviation of 0.03 in log_10_ *K_D_* (Figure 6). A second design (Rank 3) also confirmed binding with *K_D_* = 1.88 × 10^−8^ M. The rank-1 and rank-3 designs have total germline Levenshtein distances of 31 each, compared to 30 for the wild-type 1E5 Fab (Supplementary Figure S3). The top design represents a 2.8-fold improvement in binding affinity compared to the official competition winner (*K_D_* = 3.7 × 10^−10^ M, as reported by the AdaptyvBio competition organizers^1^. Our top design has low Boltz-2 confidence scores (ipSAE = 0.30, ipTM = 0.62), well below typical filtering thresholds used in design pipelines (e.g., ipTM > 0.8). AF3 assigned higher confidence to the same design (ipSAE = 0.72, ipTM = 0.87).

**Figure 6:**
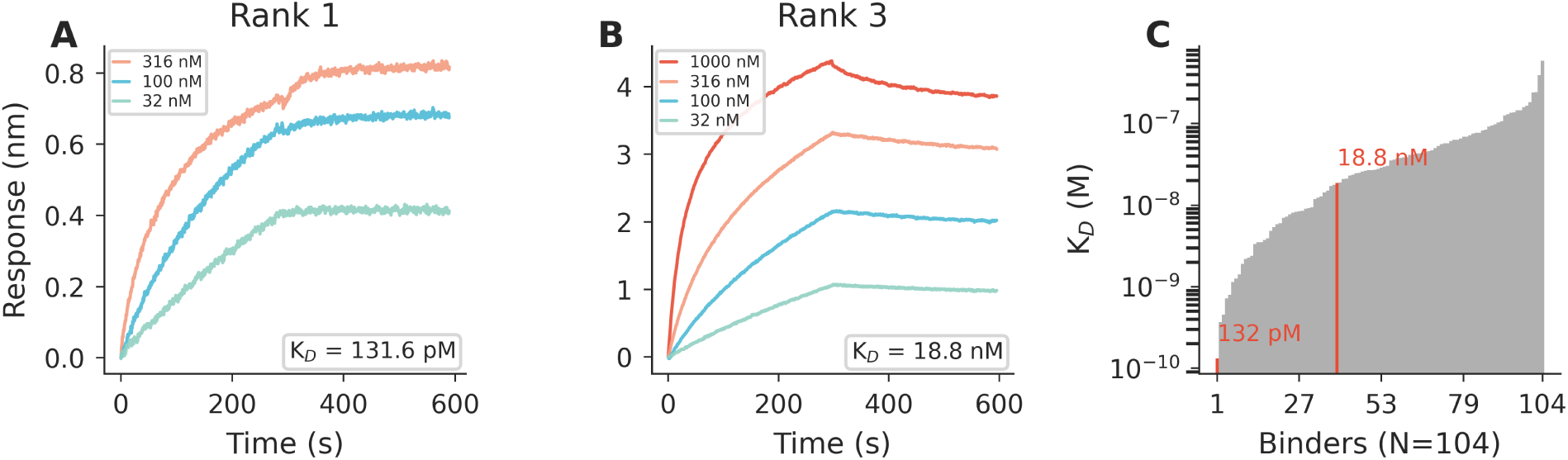
Experimental validation of ProtAff-guided Nipah virus binders. **(A)** BLI sensorgrams for the top-ranked design (Rank 1) at three analyte concentrations, yielding *K_D_* = 131.6 pM. **(B)** BLI sensorgrams for Rank 3 at four concentrations, yielding *K_D_* = 18.8 nM. Fitted curves overlaid in darker lines. **(C)** Ranked *K_D_* values across all experimentally tested competition submissions (gray), with confirmed ProtAff binders highlighted in red.

To assess whether ProtAff’s ranking favors germline reversion [58], we computed the Levenshtein distance (LD) between each of the 10,000 candidate designs and the nearest human germline V and J gene sequences, using the wild-type 1E5 Fab as the reference (Supplementary Figures S3–S5). The wild-type has a total germline LD of 30, and most mutants are more distant from germline (median total LD ≈ 34). ProtAff scores correlate weakly with total germline distance (*ρ* = 0.13) and with both naturalness scores (*ρ* = 0.08 with AntiBERTy, *ρ* = 0.12 with IgLM). The implications of these weak correlations for the independence of affinity and naturalness signals are discussed in Section 5.3.

### 4.3 Sequence-Only Models Underperform Structural Methods for Cross-Target Discrimination

We next evaluated sequence-only models on a cross-target discrimination benchmark. Smorodina et al. [5] compiled 91 experimentally resolved VHH–antigen crystal structures spanning diverse targets and reported ipTM-based cross-target discrimination results for AF3, Boltz-2, and Chai-1. We adopted their benchmark and scoring protocol for the structural predictors, and we additionally scored all 91 VHHs against each of the 91 targets using ProtAff, MINT, and CrossAffinity. For each target, the rank of the true cognate VHH among the 91 candidates was recorded; a perfect predictor would place the true binder at rank 1 for every target. We report both the top-1 hit count and the full distribution of true-binder ranks (Figure 7).

**Figure 7:**
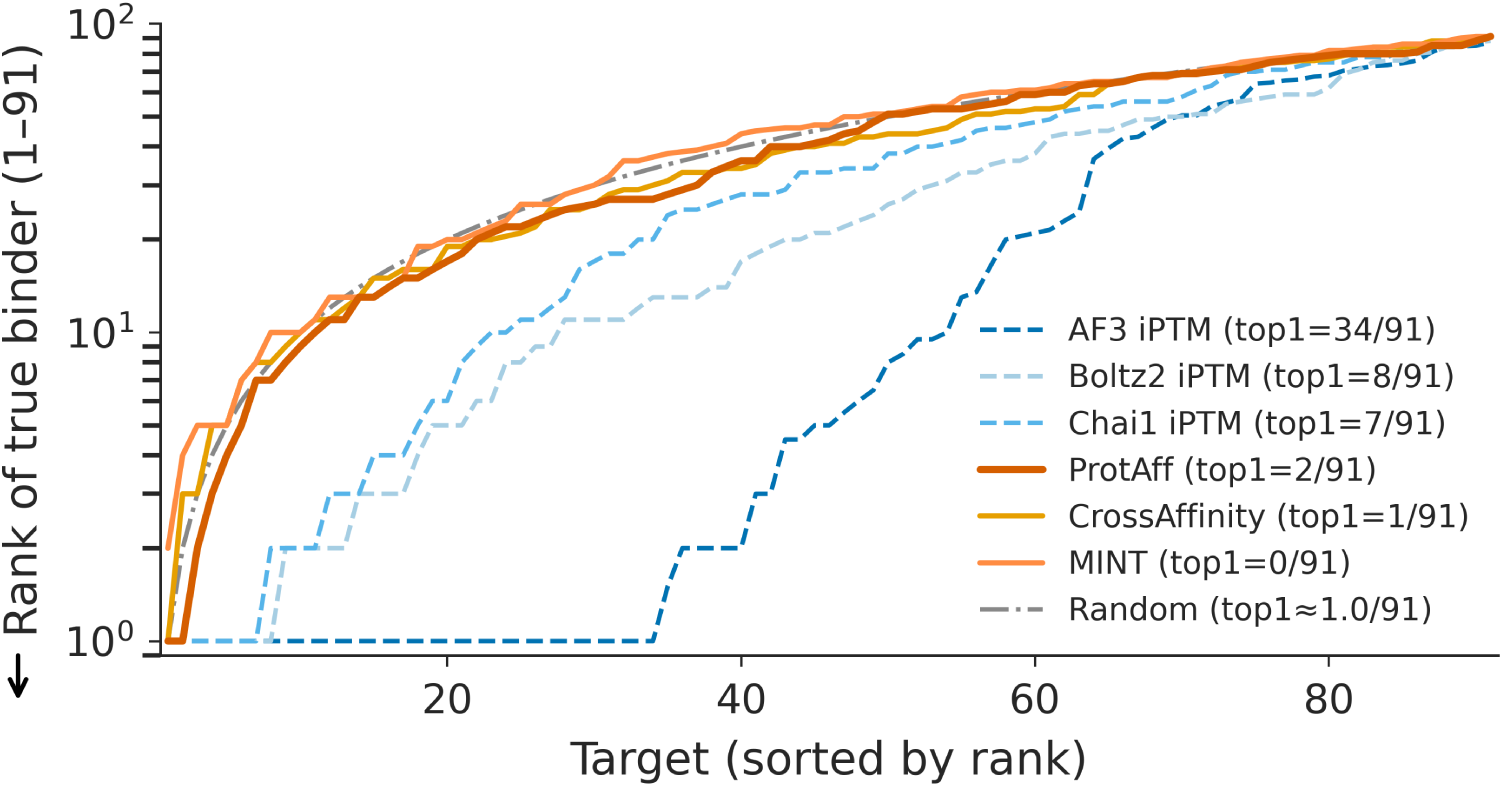
Cross-target discrimination benchmark on 91 VHH–antigen crystal structures from Smorodina et al. [5]. For each target, the 91 candidate VHHs are scored and the rank of the true cognate VHH is recorded (1 = correctly identified, 91 = worst; lower is better, indicated by the arrow on the *y*-axis). Curves show the sorted rank distribution across all 91 targets; lower curves indicate better discrimination. The gray dash-dot line shows the analytical expectation under uniform random scoring (*E*[*U*_(*i*)_] = *i* (*N*+1)/(*m*+1) for *m* = *N* = 91). Legend entries report the number of targets for which each method achieves rank 1 (top1); for the Random reference this is the expected count (≈ *m/N* = 1).

Structural methods outperform sequence-only models on this task. AF3 places the true binder at rank 1 for 34 of 91 targets, compared to 8 for Boltz-2, 7 for Chai-1, 2 for ProtAff, 1 for CrossAffinity, and 0 for MINT. The per-target rank distributions show that AF3 not only achieves the most top-1 hits but also maintains lower ranks across the full range of targets, with sequence-only models clustering together at substantially higher ranks. The separation between structural and sequence-only methods is largest at small ranks: for roughly half the targets, AF3 places the true binder at rank 5 or better, whereas no sequence-only model achieves comparable coverage below rank 10.

### 4.4 Target Ablation Reveals Limited Dependence on True Target Sequence

To interpret what the model has learned, we first analyzed the cross-attention weights and pooling weights across all 55 binder–target pairs in the EGFR benchmark. For each pair, we defined interface residues as those within 8 Å of the binding partner in the AF3-predicted structures and computed the ratio of mean attention weight at interface positions to mean attention weight at non-interface positions (Figure 8). On the binder side, the attention pooling layer assigns consistently higher weight to interface residues (median enrichment = 1.63), indicating that the model concentrates its final readout on residues involved in the interaction. On the target side, cross-attention weights show no preferential focus on interface positions (median enrichment = 1.00).

**Figure 8:**
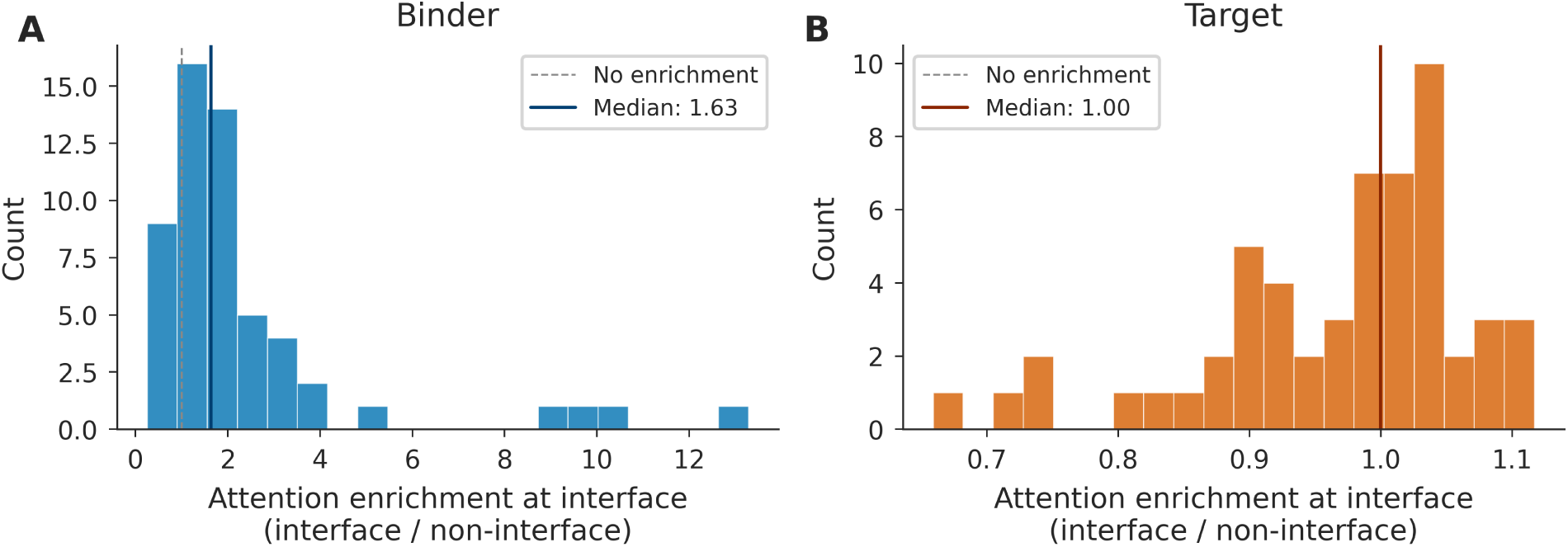
Attention enrichment at the predicted binding interface (AF3, 8 Å cutoff). **(A)** Binder pooling weight enrichment (interface / non-interface; median = 1.63). **(B)** Target cross-attention enrichment (median = 1.00).

The diffuse target-side attention, combined with the cross-target discrimination results, suggests that ProtAff makes limited use of target-specific information. To test this directly, we replaced the true EGFR target with a series of alternative targets and re-scored all 55 binders in the EGFR benchmark. We evaluated five substitutions: (i) *Scrambled EGFR*, a random permutation of the EGFR residues preserving amino acid composition (622 aa); (ii) *Most represented*, the SARS-CoV-2 Spike protein (1273 aa, 69,511 training samples), the most heavily trained-against target in ProtAff’s training set; (iii) *Similar length*, influenza hemagglutinin (519 aa, 40,854 training samples), a training target of comparable sequence length to EGFR; (iv) *Different family*, the SARS-CoV-2 Spike RBD fragment (209 aa, 53,729 training samples), a training target from a structurally unrelated protein family; and (v) *Sparse*, a cysteine-rich mini-domain (53 aa, 1 training sample), a training target with very few associated binders. For each substitution, we recomputed Spearman *ρ* and NDCG@*K* against the experimental −log *K_D_* values (Figure 9).

**Figure 9:**
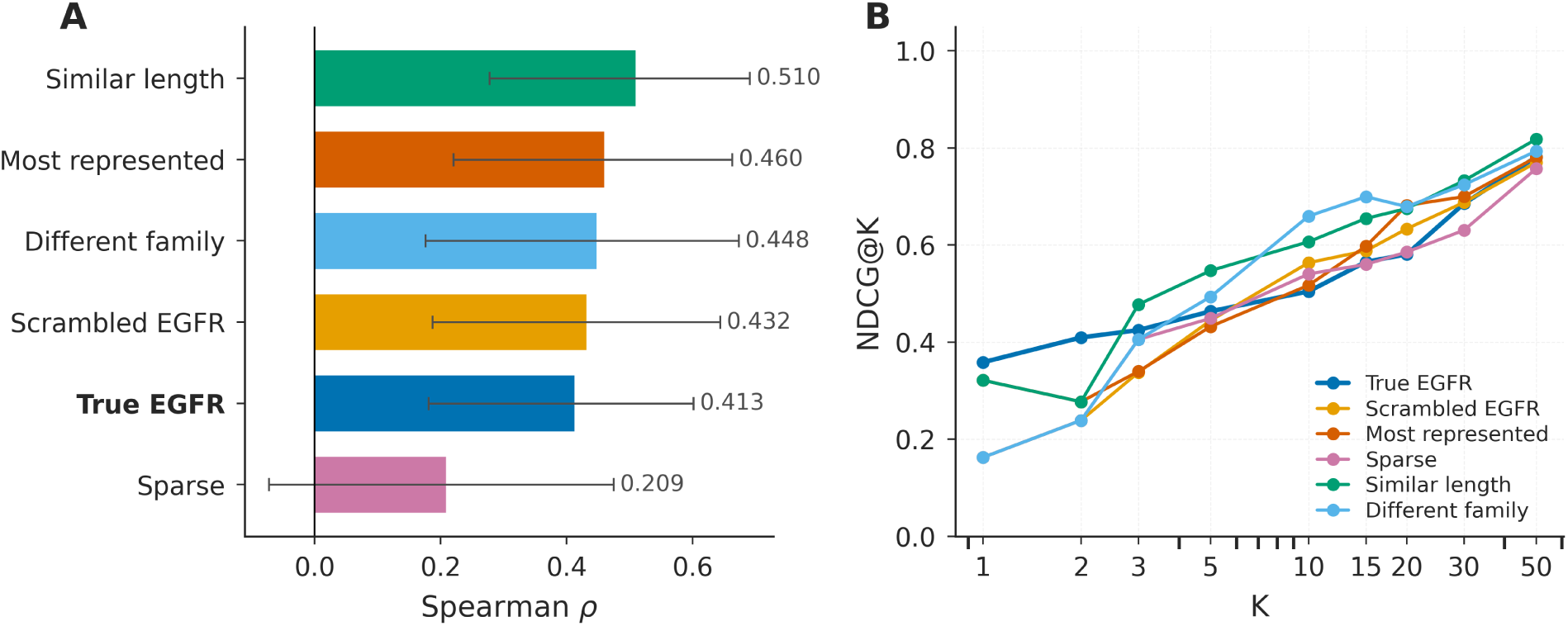
Target ablation on the AdaptyvBio EGFR benchmark (*N* = 55). For each of the 55 binders, the true EGFR target is replaced with an alternative target before scoring. **(A)** Spearman *ρ* between predicted affinity and experimental −log *K_D_* under each target substitution, with error bars from 1,000 bootstrap resamples. **(B)** NDCG@*K* curves for the same conditions.

Four of the five substitutions yielded Spearman correlations indistinguishable from or higher than the true EGFR target (True EGFR *ρ* = 0.413; Scrambled EGFR *ρ* = 0.432; Different family *ρ* = 0.448; Most represented *ρ* = 0.460; Similar length *ρ* = 0.510). Only the Sparse target produced a clear drop (*ρ* = 0.209). NDCG@*K* curves show a similar pattern, with most substitutions tracking or slightly exceeding True EGFR across the full range of *K*. The Sparse substitution trails at small *K* but converges with the other conditions by *K* = 20. The relationship between substitution target, training set representation, and ranking performance is examined in Section 5.5.

### 4.5 Biophysical Property Correlations Suggest Confounding with Binder Composition

The weak dependence on the true target sequence raises the question of what binder-side signals drive ProtAff’s predictions. We computed Spearman correlations between four biophysical properties of the binder (sequence length, mean hydrophobicity, net charge at pH 7, and isoelectric point) and two quantities: the ProtAff predicted affinity score, and the experimental −log *K_D_* (Figure 10).

**Figure 10:**
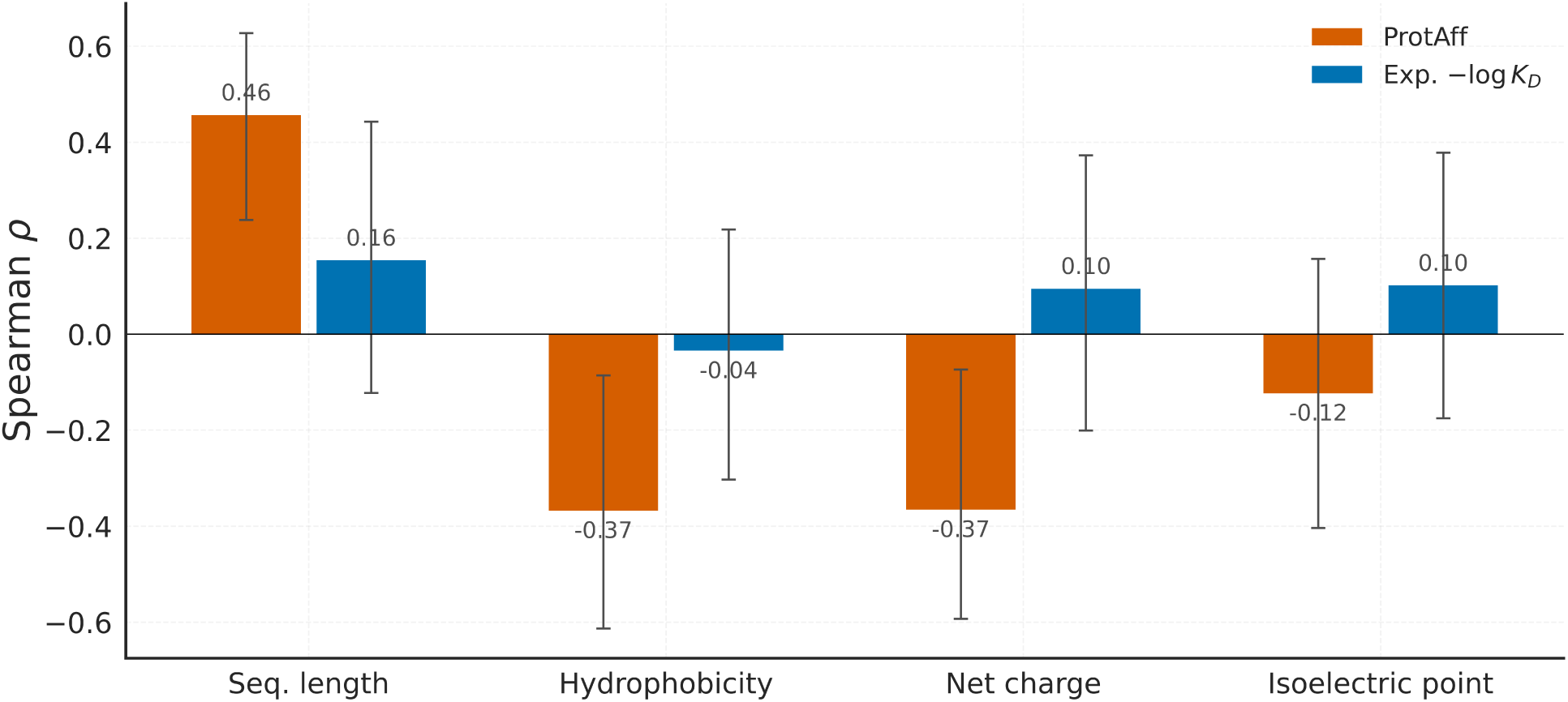
Spearman correlation between binder biophysical properties and both the ProtAff predicted affinity (orange) and the experimental −log *K_D_* (blue) on the AdaptyvBio EGFR benchmark (*N* = 55). Error bars show 95% confidence intervals from 1,000 bootstrap resamples.

Three of the four properties correlate more strongly with the predicted score than with the experimental measurement. Sequence length shows the largest discrepancy: predicted affinity correlates with length at *ρ* = 0.46 (longer binders receive higher predicted affinity scores), whereas the experimental correlation is only *ρ* = 0.16. Hydrophobicity correlates at *ρ* = −0.37 with predicted affinity but only *ρ* = −0.04 with experimental −log *K_D_*. Net charge shows a similar magnitude (*ρ* = −0.37 for predicted, *ρ* = 0.10 for experimental), with the two correlations running in opposite directions. Isoelectric point shows a weaker version of the same effect (*ρ* = −0.12 for predicted, *ρ* = 0.10 for experimental). The origins and implications of these discrepancies are examined in Section 5.5.

We repeated the correlation analysis using the predicted scores from each of the five target-substitution conditions described in Section 4.4 (Supplementary Figure S7 and Supplementary Table S4). Sequence length correlates positively with the predicted score in all six conditions (*ρ* ≈ 0.04 to 0.62), reaching significance in five (*p <* 0.05), including the scrambled control (*ρ* = 0.43) where real binding signal is destroyed but binder composition is unchanged. Net charge and isoelectric point show consistent positive correlations across five of six conditions (*ρ* ≈ 0.3 to 0.5), with the true EGFR condition as the sign-flipped outlier. Hydrophobicity is the least stable property, with signs flipping across conditions and most confidence intervals crossing zero. That the sequence-length and charge correlations persist at full magnitude under scrambled EGFR confirms that these biases are read from the binder sequence itself rather than inferred from target context. None of the four features correlates significantly with the experimental −log *K_D_* (all |*ρ*| *<* 0.16, *p >* 0.25).

## 5 Discussion

### 5.1 Structure Prediction Versus Affinity Prediction

On the AdaptyvBio EGFR benchmark, all AF3 and Boltz-2 structural confidence metrics showed negative or near-zero Spearman correlations with experimental binding affinity (Table 1, Supplementary Table S3), while ProtAff achieved *ρ* = 0.413 using sequence alone. In the Nipah competition, our top-ranked design has low Boltz-2 confidence (Section 4.2) despite binding at 131.6 pM, and it was discarded under the competition’s standard filtering thresholds. These results indicate that structural confidence scores, which are trained on experimentally resolved complexes and reflect pose plausibility rather than binding energetics, are effective for rejecting misfolded interfaces but not for ranking affinity among designs that all produce plausible poses. ipSAE (min) illustrates this directly: it achieves NDCG@5 of 0.666 despite Spearman *ρ* = 0.054 because the strongest binders likely form the most well-resolved interfaces, but among candidates that all produce plausible complexes, the confidence scores cannot distinguish binding strength. King et al. [59] reached the same conclusion from an independent direction: after adapting the Boltz-2 ligand affinity head for protein–protein affinity prediction on the TCR3d [60] and PPB-affinity [25] datasets, they found that fine-tuned ESM-2 sequence embeddings outperformed Boltz-2 structural embeddings, with combining the two modalities yielding only marginal gains over sequence alone.

### 5.2 Heterogeneous Data and the Ranking Loss

The margin ranking loss (described in Section 2.3) is central to training on heterogeneous data because it sidesteps the residual calibration differences between assay types. Combined with inter-target pairing, it encourages the model to learn a globally coherent affinity ranking without requiring compatible measurement scales. The severe target imbalance (Figure 1) poses an additional challenge: without correction, the model would overfit to a handful of well-characterized targets. Cluster-balanced pair sampling reduces this dominance, and the group-based validation split ensures that validation targets are entirely unseen during training.

The two ProtAff configurations illustrate a second consequence of the loss function choice. The weighted variant, which scales margin penalties by the affinity gap between paired samples, prioritizes correct ordering among the strongest binders at the cost of full-list ranking accuracy. The choice between the two variants therefore depends on whether the downstream application prioritizes selecting a small number of top candidates for experimental validation or requires accurate ordering across the full ranked list.

### 5.3 Nipah Case Study

The Nipah virus competition results illustrate both the utility and the limitations of ProtAff in a practical design pipeline. Our top design achieved sub-nanomolar binding stronger than the competition winner (Figure 6), validating ProtAff’s ability to rank binding affinity among candidate designs. The disagreement between Boltz-2 and AF3 on this design (Section 4.2) underscores that structural confidence scores can be unreliable for individual predictions even when they perform well on average, and that an orthogonal affinity signal may help identify strong binders missed by structural filtering alone.

The weak correlations between ProtAff scores and both germline distance and naturalness metrics (Section 4.2) indicate that Stage 3 affinity ranking captures features distinct from Stage 2 naturalness filtering, ruling out the concern that ProtAff favors germline-proximal sequences [58]. However, with only 5 designs tested, the 60% failure rate (3 of 5 did not express) is too small a sample to determine whether the pipeline systematically selects for poorly expressing sequences or whether these failures reflect the typical expression rate for de novo designs. ProtAff does not model protein expressibility or stability, and integrating expressibility prediction into the pipeline could improve the overall hit rate.

### 5.4 Cross-Target Discrimination

While structural confidence scores fail at within-target affinity ranking, our cross-target discrimination benchmark (Figure 7) shows the reverse: AF3 places the true cognate VHH at rank 1 for 34 of 91 targets, compared to at most 2 for any sequence-only method. Structural methods retain a clear advantage when the task requires identifying which target a given binder recognizes, consistent with cross-target discrimination being a structural compatibility problem that sequence-only models cannot fully recover.

AF3’s strong per-target ranking here may seem inconsistent with Smorodina et al. [5], who reported high false positive rates for ipTM-based cognate discrimination. Smorodina et al. evaluate precision-recall across all 91 targets jointly, sweeping a threshold over the pooled score distribution; because many non-cognate pairs receive high ipTM, precision is low at every threshold that recovers a reasonable fraction of true cognates. Our protocol ranks candidates within each target independently, where only the relative ordering matters. AF3’s scores are informative enough to place the true binder near the top within many individual targets, even though the pooled score distributions do not separate cognate from non-cognate pairs cleanly.

### 5.5 Target Dependence and Biophysical Confounds

Despite the architectural design of the cross-attention module to condition predictions on the target sequence, the attention enrichment and target ablation analyses (Figures 8 and 9) indicate that within-target ranking on the EGFR benchmark is largely driven by binder-side features. The model concentrates pooling on binder interface residues but distributes cross-attention diffusely across the target, and replacing the true EGFR sequence with unrelated targets preserves or even improves ranking performance.

Since four of five substitutions are unrelated to EGFR in sequence and function, the ranking cannot depend on target-specific interaction modeling. The Sparse condition’s degradation is consistent with this: its single training sample means the model has rarely seen any binder scored against it, limiting the quality of the conditioning signal. The biophysical property analysis identifies what the model relies on instead: the training distribution is dominated by antibody variable domains and computationally designed mini-binders with characteristic length, hydrophobicity, and charge profiles, and predicted affinity correlates with these binder properties more strongly than experimental affinity does (Figure 10). These correlations persist under scrambled EGFR where real binding signal is destroyed, confirming that the model treats binder-level compositional profiles as affinity cues. The cross-target correlation analysis (Supplementary Figure S7 and Supplementary Table S4) reinforces this interpretation. Swapping the target preserves the overall ranking of binders (Spearman *ρ* with experimental affinity is maintained or improved in four of five conditions), but changes which binder features correlate with the predicted score: sequence-length bias persists across five of six conditions, while charge-related correlations flip sign depending on the conditioning target. Since the binder properties are fixed across conditions, the target embedding modulates which binder features the model weights as favorable rather than encoding interaction-specific binding physics.

These findings do not invalidate ProtAff’s practical utility but clarify what the model captures and where it fails. Mitigating target-independence may require training objectives that explicitly penalize target-agnostic scoring, such as contrastive losses using target-shuffled negatives, or training data with more balanced binder property distributions across diverse target families.

### 5.6 Limitations

Both the margin ranking loss and LoRA fine-tuning contribute to performance (Table 1), but we have not systematically varied other architectural hyperparameters (number of cross-attention layers, LoRA rank, depth of LoRA application), so the current configuration may not be optimal. The attention enrichment analysis uses AF3-predicted structures rather than experimental crystal structures, introducing potential circularity given that these predictors vary in accuracy on this benchmark. Whether the binder-level compositional biases identified in the target ablation and biophysical property analyses (Section 5.5) persist on benchmarks with more diverse binder scaffolds remains untested.

As demonstrated by the cross-target discrimination benchmark (Figure 7), sequence-only models perform poorly at distinguishing cognate from non-cognate pairings, with structural methods holding a substantial advantage. Cross-target specificity might be improved by incorporating non-cognate binder–target pairs as negative training examples, though curating reliable negatives without introducing false labels remains an open challenge.

The evaluation is limited in scope. The EGFR benchmark comprises only *N* = 55 binders against a single target, limiting statistical power and the ability to assess generalization across diverse target families. Evaluating ProtAff on additional held-out targets with verified no training overlap would provide a more rigorous assessment of cross-target generalization. The training data are dominated by antibody–antigen interactions; performance on other PPI classes (e.g., enzyme–inhibitor, receptor–ligand) with sparse training data remains unknown.

## 6 Conclusion

ProtAff demonstrates that sequence-only affinity prediction, combining LoRA-finetuned ESM-2 with cross-attention and margin ranking loss, outperforms both structural confidence scores and existing ML-based predictors for within-target ranking (*ρ* = 0.413, NDCG@5 = 0.758 for the weighted variant, on the AdaptyvBio EGFR benchmark). A pipeline incorporating ProtAff produced a sub-nanomolar Nipah virus binder (131.6 pM) that exceeded the competition winner by 2.8-fold. However, target ablation and biophysical property analyses show that ProtAff’s ranking on the EGFR benchmark correlates with binder-level compositional features rather than target-conditioned interaction modeling, and structural methods retain clear advantages for cross-target discrimination (AF3: 34/91 top-1 hits versus ProtAff: 2/91). Future work will focus on training objectives that enforce target-dependent scoring, integration of expressibility prediction, and multi-task learning across diverse target families.

## Acknowledgments

This work was supported by National Institutes of Health grant R35-GM141881, the National Science Foundation (NSF) Graduate Research Fellowship Program (MC), and AstraZeneca. Computational resources were provided by the Advanced Research Computing at Hopkins (ARCH).

## Data Availability Statement

The inference code, model weights, and test set are available at https://github.com/Graylab/ProtAff.

## Supporting Information

### Training Data Composition

**Table S1:**
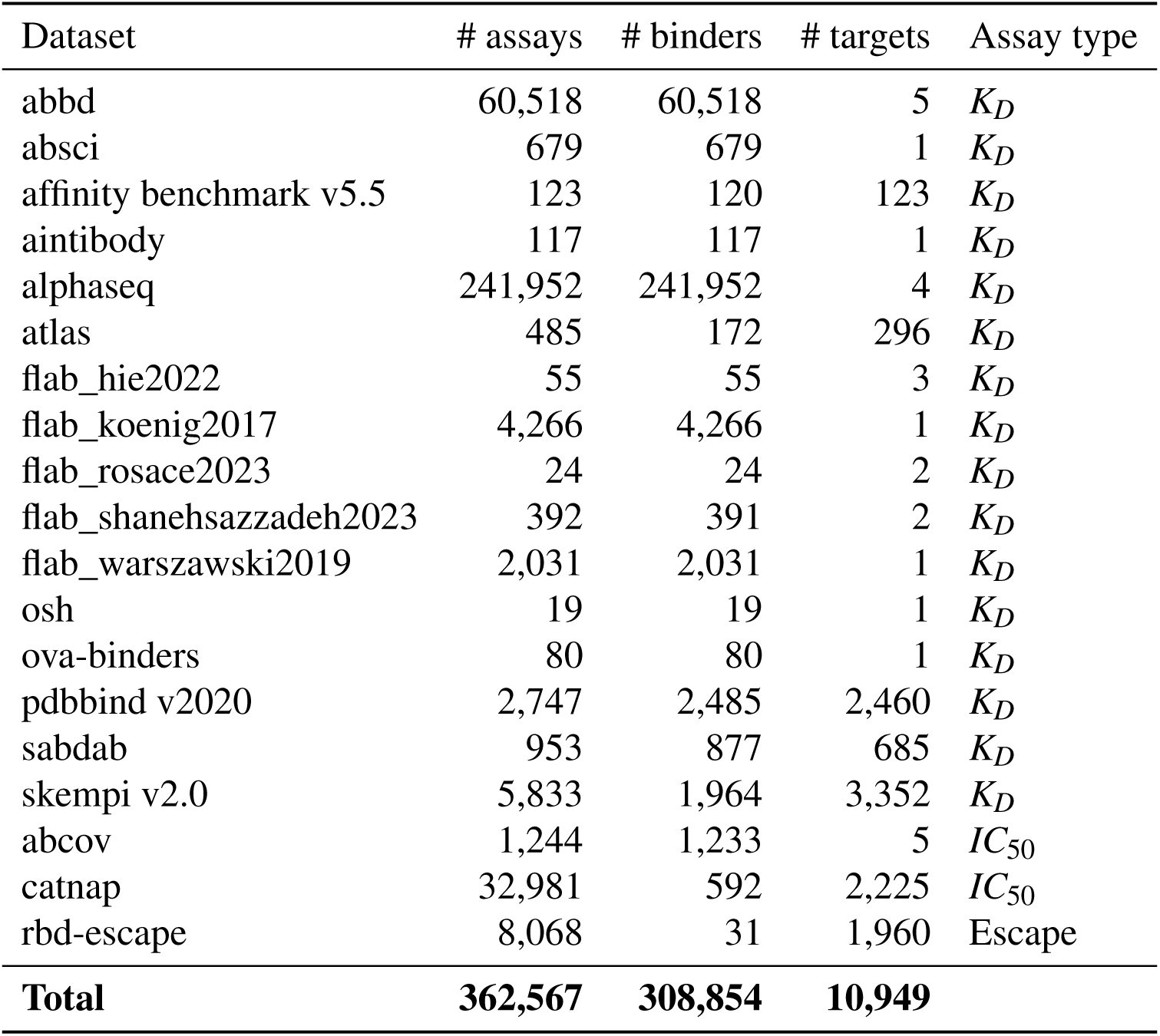
Composition of the ProtAff training corpus by source. All datasets were accessed through the AbRank [14], ASD [24], or PPB-Affinity [25] aggregated databases.

### Hyperparameters

**Table S2:**
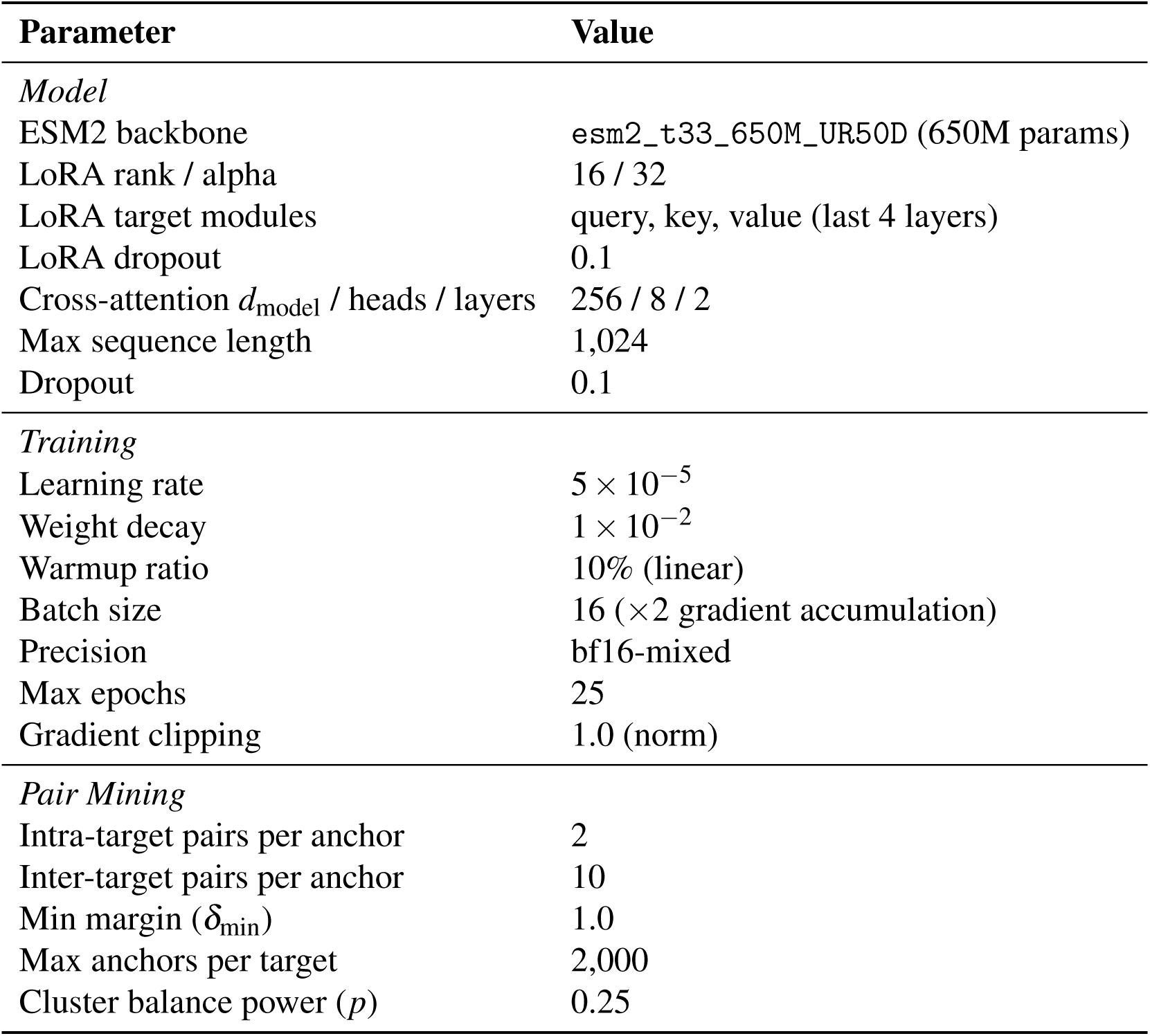
Model and training hyperparameters.

### Structural Metric Aggregation Comparison

**Table S3:** Evaluation of structural confidence metrics across aggregation methods for both AF3 and Boltz-2 on the AdaptyvBio EGFR benchmark (*N* = 55). Values: mean± std over 1,000 bootstrap resamples.

| Source | Metric (agg) | Spearman $\rho$ | Pearson $R$ | NDCG@5 |
| --- | --- | --- | --- | --- |
| **AF3** |  |  |  |  |
| AF3 | ipSAE (max) | -0.112±0.151 | -0.102±0.158 | 0.422±0.122 |
| | ipSAE (min) | 0.054±0.156 | 0.013±0.164 | 0.666±0.202 |
| | ipSAE (mean) | -0.064±0.154 | -0.061±0.160 | 0.747±0.181 |
| | ipSAE (median) | -0.114±0.151 | -0.078±0.157 | 0.422±0.121 |
| | ipTM (max) | -0.151±0.144 | -0.174±0.154 | 0.311±0.105 |
| | ipTM (min) | -0.141±0.144 | -0.137±0.157 | 0.311±0.105 |
| | ipTM (mean) | -0.142±0.145 | -0.158±0.156 | 0.311±0.105 |
| | ipTM (median) | -0.113±0.149 | -0.155±0.157 | 0.311±0.105 |
| | pDockQ (max) | -0.121±0.152 | -0.052±0.159 | 0.185±0.074 |
| | pDockQ (min) | -0.102±0.152 | -0.083±0.162 | 0.274±0.084 |
| | pDockQ (mean) | -0.117±0.149 | -0.077±0.155 | 0.187±0.094 |
| | pDockQ (median) | -0.125±0.147 | -0.093±0.147 | 0.181±0.096 |
| | pDockQ2 (max) | -0.110±0.155 | -0.083±0.158 | 0.705±0.174 |
| | pDockQ2 (min) | -0.158±0.156 | -0.098±0.156 | 0.694±0.179 |
| | pDockQ2 (mean) | -0.165±0.157 | -0.091±0.159 | 0.710±0.193 |
| | pDockQ2 (median) | -0.138±0.159 | -0.084±0.158 | 0.726±0.181 |
| | LIS (max) | -0.187±0.142 | -0.125±0.153 | 0.297±0.074 |
| | LIS (min) | -0.094±0.144 | -0.076±0.152 | 0.265±0.058 |

Table S3 – continued from previous page
| Source | Metric (agg) | Spearman $\rho$ | Pearson $R$ | NDCG@5 |
| --- | --- | --- | --- | --- |
|  | LIS (mean) | -0.130±0.142 | -0.105±0.152 | 0.231±0.043 |
|  | LIS (median) | -0.141±0.141 | -0.110±0.152 | 0.231±0.044 |
| **Boltz-2** |  |  |  |  |
| Boltz-2 | ipSAE (max) | -0.181±0.145 | 0.029±0.135 | 0.262±0.056 |
|  | ipSAE (min) | -0.046±0.146 | -0.068±0.150 | 0.218±0.055 |
|  | ipSAE (mean) | -0.179±0.143 | -0.045±0.142 | 0.250±0.044 |
|  | ipSAE (median) | -0.194±0.142 | -0.063±0.142 | 0.262±0.056 |
|  | ipTM (max) | -0.150±0.137 | 0.004±0.131 | 0.240±0.047 |
|  | ipTM (min) | -0.243±0.132 | -0.162±0.128 | 0.271±0.055 |
|  | ipTM (mean) | -0.176±0.135 | -0.090±0.128 | 0.231±0.044 |
|  | ipTM (median) | -0.170±0.134 | -0.087±0.123 | 0.252±0.041 |
|  | pDockQ (max) | -0.040±0.142 | -0.120±0.150 | 0.204±0.085 |
|  | pDockQ (min) | -0.156±0.144 | -0.124±0.146 | 0.223±0.073 |
|  | pDockQ (mean) | -0.111±0.147 | -0.120±0.150 | 0.206±0.081 |
|  | pDockQ (median) | -0.120±0.146 | -0.112±0.158 | 0.206±0.082 |
|  | pDockQ2 (max) | -0.100±0.142 | -0.101±0.124 | 0.239±0.073 |
|  | pDockQ2 (min) | -0.157±0.140 | -0.136±0.120 | 0.247±0.072 |
|  | pDockQ2 (mean) | -0.108±0.141 | -0.138±0.125 | 0.239±0.080 |
|  | pDockQ2 (median) | -0.162±0.141 | -0.173±0.126 | 0.247±0.068 |
|  | LIS (max) | -0.188±0.148 | -0.026±0.141 | 0.323±0.064 |
|  | LIS (min) | -0.226±0.142 | -0.172±0.144 | 0.276±0.043 |
|  | LIS (mean) | -0.222±0.144 | -0.127±0.143 | 0.278±0.047 |
|  | LIS (median) | -0.234±0.142 | -0.131±0.143 | 0.278±0.047 |

### Sequence Similarity Between AbRank Training and Test Splits

We examined the sequence composition of the AbRank benchmark [14] using the publicly released splits^2^. For each test-set example, we computed the maximum sequence identity of its binder and target to entries in the training set. Figure S1 shows the resulting joint distribution. Most test examples share high sequence identity with training entries on both axes, with relatively few falling into the novel-binder, novel-target quadrant. Performance on the standard split therefore primarily reflects predictions on sequences closely related to training, which is useful context when interpreting reported metrics or designing evaluations targeting novel binders and targets.

**Figure S1:**
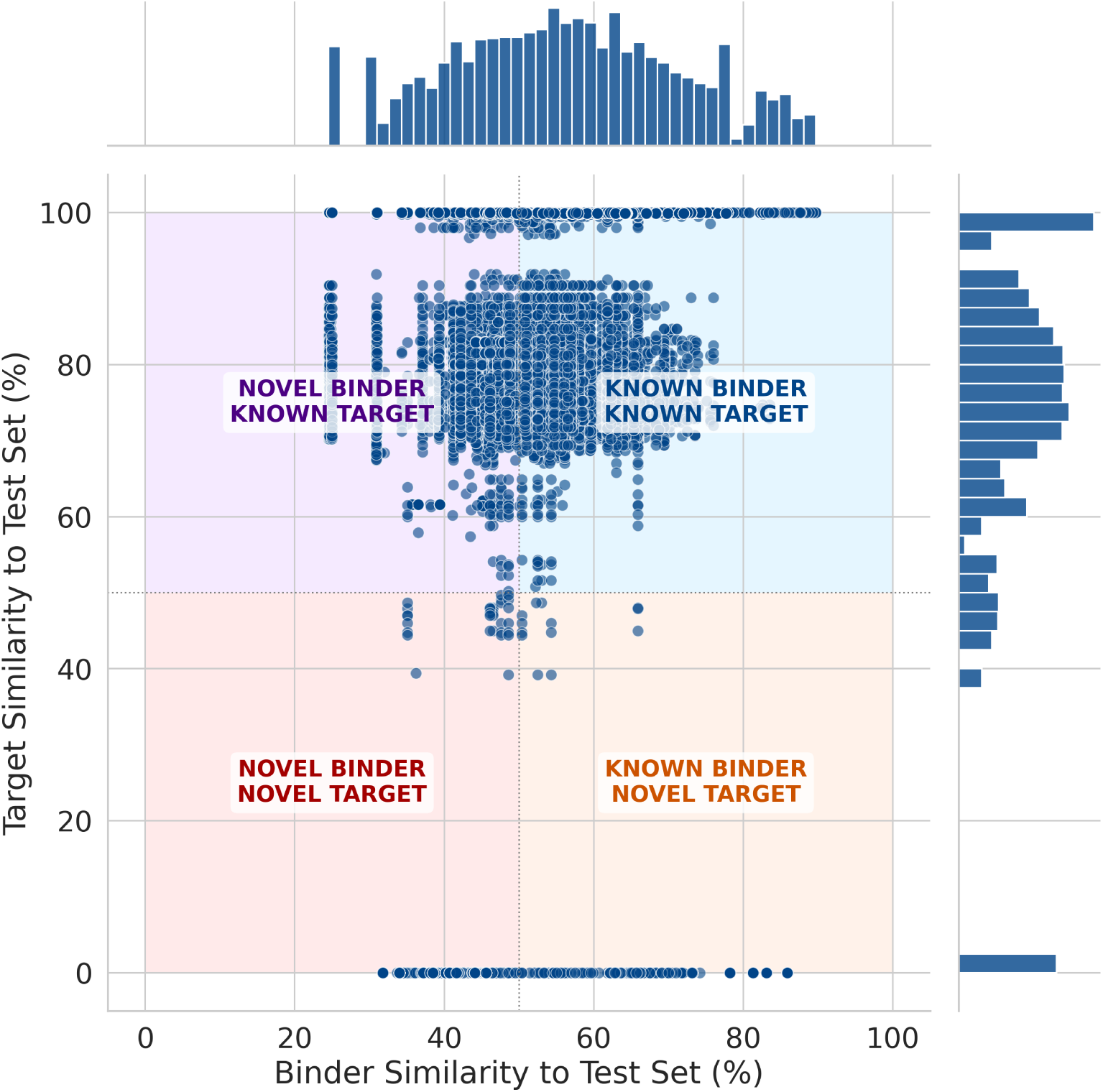
Sequence similarity between AbRank training and test splits. Each point represents a test-set example plotted by the maximum sequence identity of its binder (x-axis) and target (y-axis) to the training set. The four quadrants partition test examples by whether the binder and target are novel (less than 50% identity) or known (greater than 50% identity) relative to training. Marginal histograms show the distribution along each axis.

### Design Pipeline

**Figure S2:**
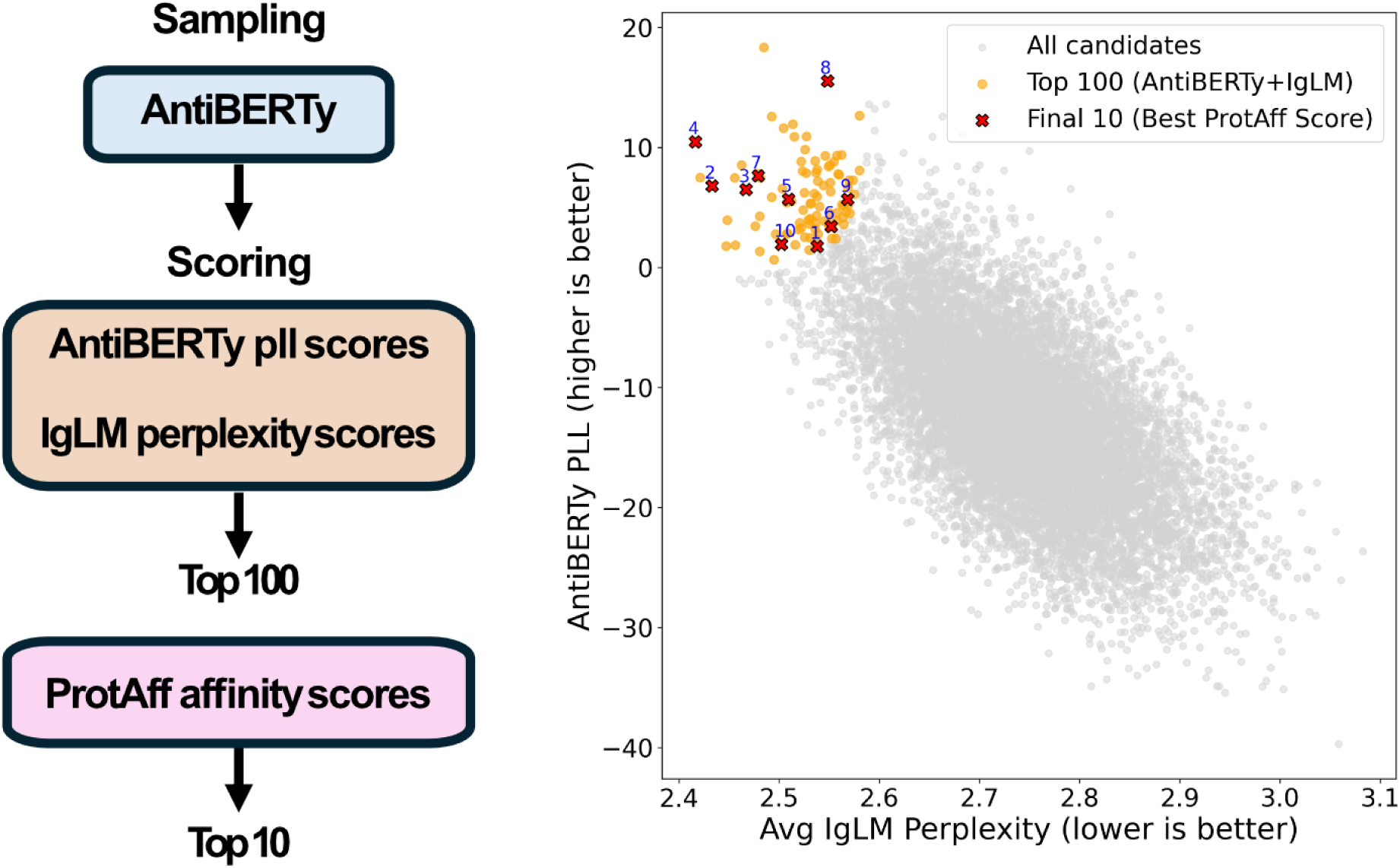
Three-stage binder design pipeline for the Nipah virus competition. **Stage 1 (sampling):** AntiBERTy masked language modeling identifies mutable positions in the 1E5 Fab [55] (PDB: 8K0C) and generates 10,000 candidate variants by stochastic mutation sampling. **Stage 2 (naturalness filtering):** Candidates are scored by combined AntiBERTy pseudo-log-likelihood and IgLM perplexity, retaining the top 100 designs with the most natural-like sequences. **Stage 3 (affinity ranking):** ProtAff scores each candidate against Nipah glycoprotein G (PDB: 2VSM) to select the top 5 designs for experimental validation by biolayer interferometry.

### Germline Distance Analysis

**Figure S3:**
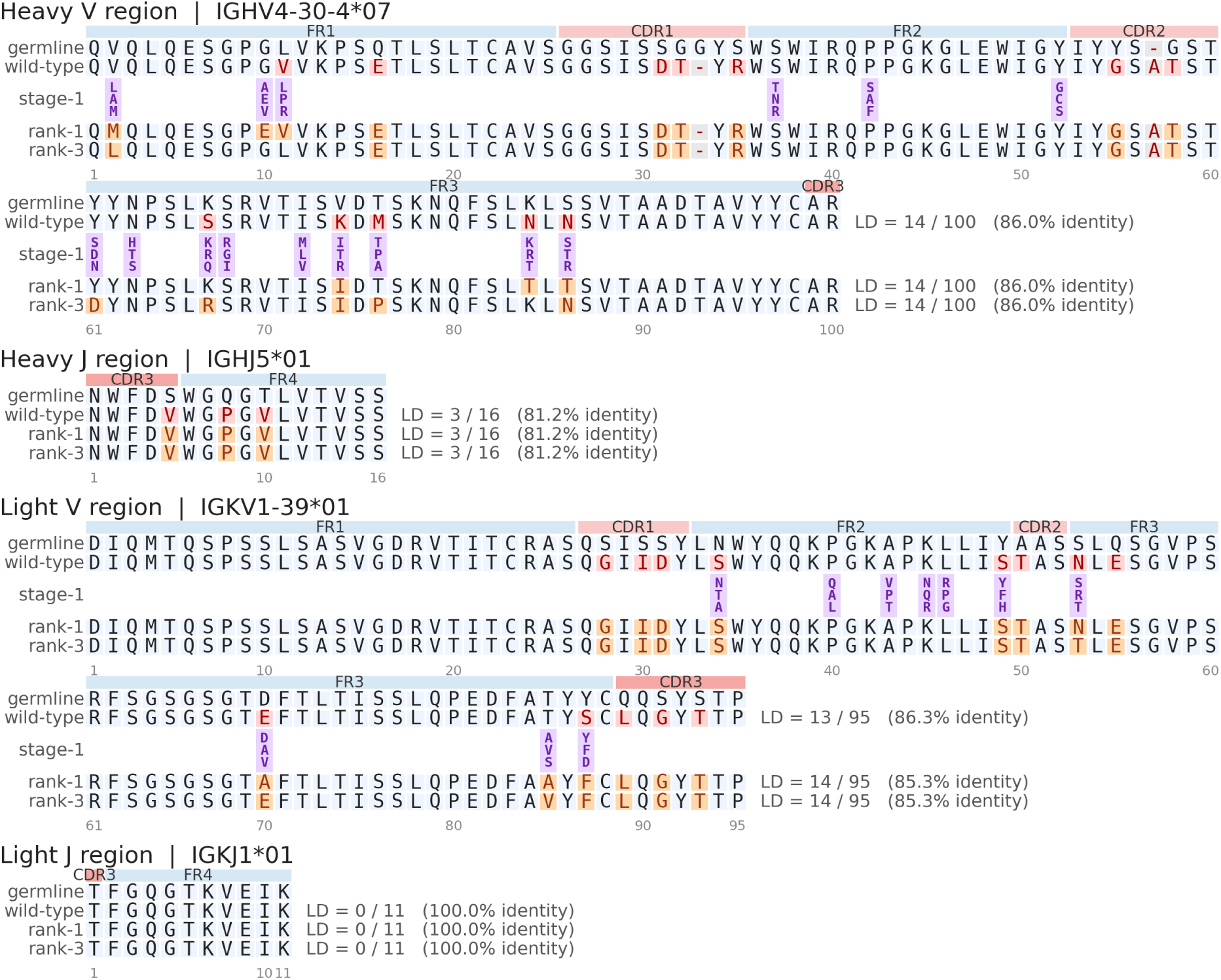
Alignment of wild-type 1E5 Fab (PDB: 8K0C) heavy and light chain sequences to their nearest human germline V and J genes, shown alongside the top-ranked designed candidates (rank-1 and rank-3). Each panel shows a gapped alignment with cells colored by germline match: positions matching the germline are highlighted in blue, wild-type residues that diverge from the germline are highlighted in red, and candidate residues that diverge from the germline are highlighted in orange. The “stage-1” row, inserted between the wild-type and candidate rows, marks the mutation-tolerant positions identified by AntiBERTy in the sampling stage; each purple cell stacks the top-ranked candidate residues (by delta-logit) considered at that position. IMGT framework (blue) and CDR (pink) regions are annotated above each alignment. Levenshtein distances and percent identity to the germline are reported per sequence row.

**Figure S4:**
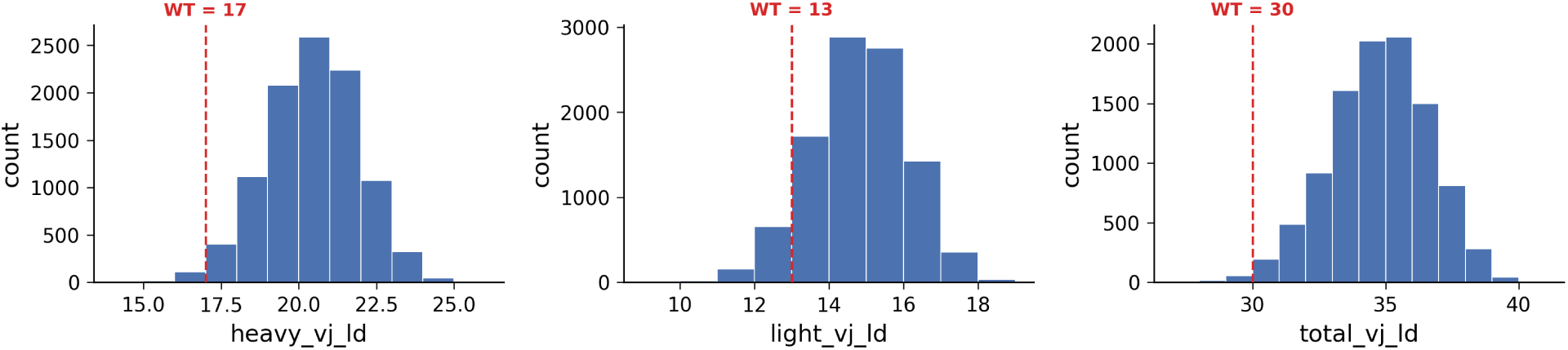
Distribution of germline Levenshtein distances across 10,000 Nipah binder candidate designs. Histograms show heavy chain V+J distance (left), light chain V+J distance (center), and total V+J distance (right). Red dashed lines mark the wild-type distances.

**Figure S5:**
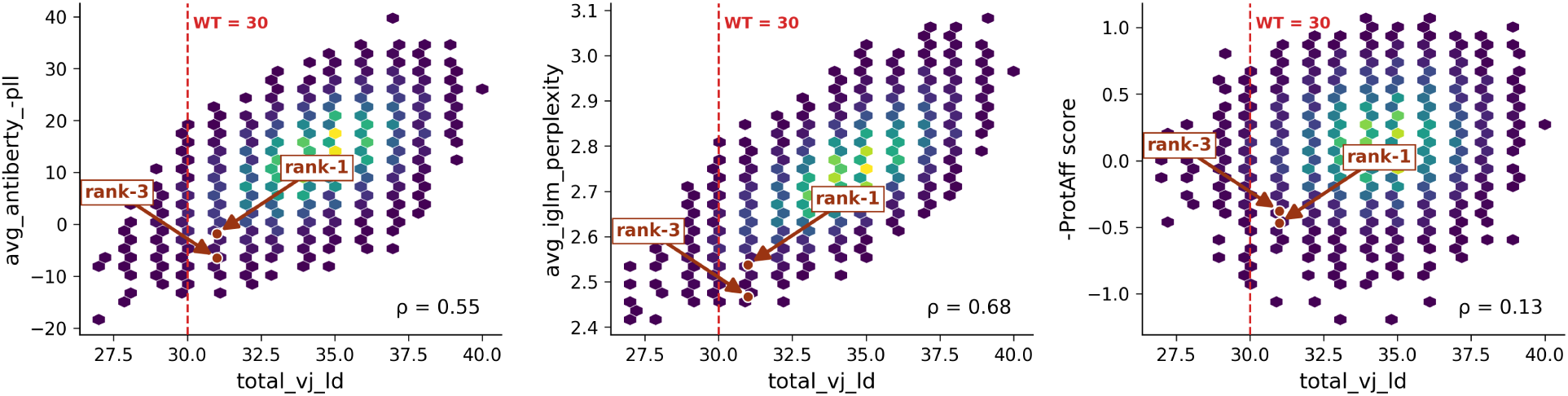
Relationship between germline distance and pipeline scoring metrics for 10,000 Nipah binder designs. Hexbin scatter plots (lighter (yellow) = higher density of candidates, and darker (purple/blue) = lower density) show total V+J germline Levenshtein distance versus AntiBERTy pseudo-log-likelihood (left), IgLM perplexity (center), and negated ProtAff score (right). Red dashed line marks wild-type total germline distance (LD = 30); orange markers and arrows indicate the rank-1 and rank-3 (both at LD = 31). Spearman *ρ* is shown in each panel.

**Figure S6:**
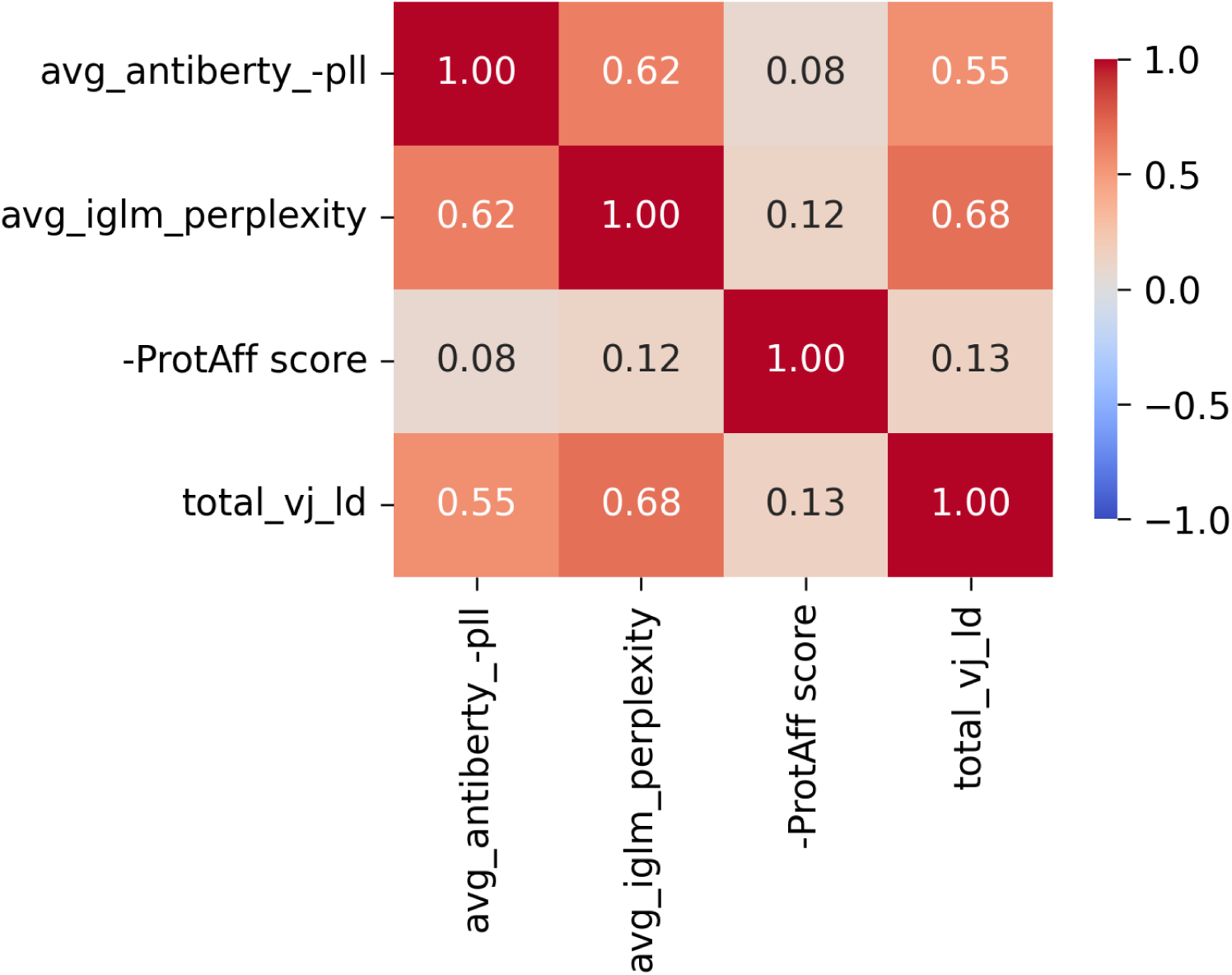
**Spearman correlation heatmap between pipeline scoring metrics and total germline distance for the 10,000 Nipah binder designs.**

### Cross-Target Biophysical Correlation Analysis

**Figure S7:**
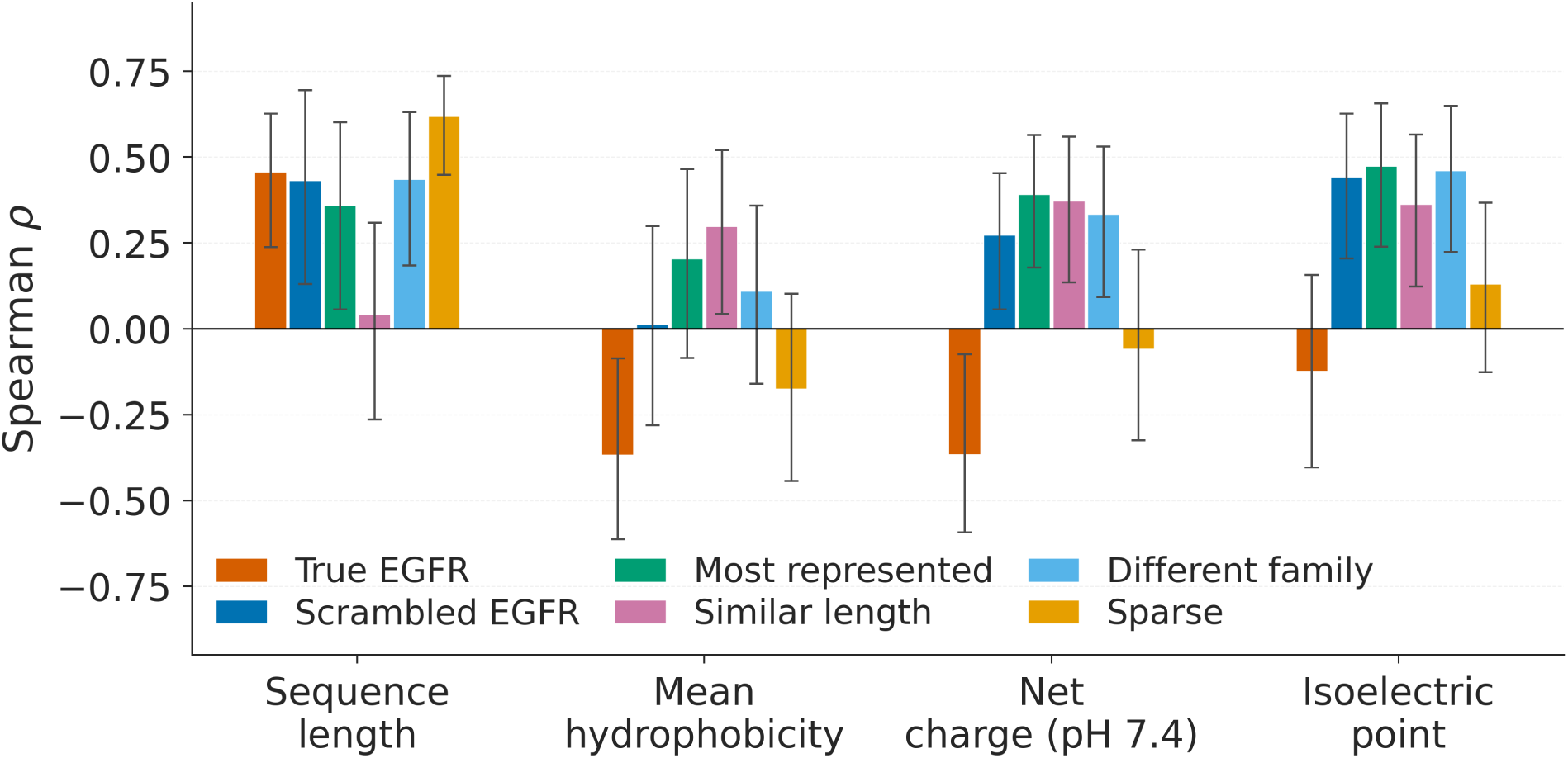
Spearman correlation between binder biophysical properties and ProtAff predicted affinity under each target-substitution condition (*N* = 55 AdaptyvBio EGFR binders). Error bars show 95% confidence intervals from 1,000 bootstrap resamples.

**Table S4:** Spearman *ρ* between binder biophysical properties and ProtAff predicted affinity under each target-substitution condition (*N* = 55). Asterisks denote *p <* 0.05.

| Condition | Sequence length | Hydrophobicity | Net charge (pH 7.4) | Isoelectric point |
| --- | --- | --- | --- | --- |
| True EGFR | −0.46* | +0.37* | +0.37* | +0.12 |
| Scrambled EGFR | −0.43* | −0.01 | −0.27* | −0.44* |
| Most represented | −0.36* | −0.20 | −0.39* | −0.48* |
| Similar length | −0.04 | −0.30* | −0.37* | −0.36* |
| Different family | −0.44* | −0.11 | −0.33* | −0.46* |
| Sparse | −0.62* | +0.18 | +0.06 | −0.13 |

## Footnotes

1 https://proteinbase.com/collections/nipah-binder-competition-results

2 https://www.kaggle.com/datasets/aurlienplissier/abrank?select=Benchmarks

